# Linking neuronal and hemodynamic network signatures in the resting human brain

**DOI:** 10.1101/2022.08.28.505586

**Authors:** Adham Elshahabi, Silke Ethofer, Holger Lerche, Daniel van de Velden, Hans Wehrl, Christian la Fougère, Christoph Braun, Niels K. Focke

**Affiliations:** Department of Neurology and Epileptology, Hertie Institute for Clinical Brain Research, University Hospital Tübingen, D-72076 Tübingen, Germany; Werner Reichardt Centre for Integrative Neuroscience, D-72076 Tübingen, Germany; MEG Center, University of Tübingen, D-72076 Tübingen, Germany; International Max Planck Research School for Cognitive and Systems Neuroscience, D-72076 Tübingen, Germany; Department of Nuclear Medicine and Clinical Molecular Imaging, University Hospital Tübingen, D-72076 Tübingen, Germany; CIMeC, Center for Mind/Brain Sciences, University of Trento, I-38123 Trento, Italy; Clinic for Neurology, University Medical Center Göttingen, D-37075 Göttingen, Germany; Werner Siemens Imaging Center, Department of Preclinical Imaging and Radiopharmacy, Eberhard Karls University, D-72076 Tuebingen, Germany

**Keywords:** EEG-fMRI, blood-oxygenation-level-dependent contrast, resting state networks, hemodynamic response function

## Abstract

Despite several studies investigating the relationship between blood-oxygen-level-dependent functional MRI (BOLD-fMRI) and neuroelectric activity, our understanding is rather incomplete. For instance, the canonical hemodynamic response function (HRF) is commonly used, regardless of brain region, frequency of electric activity and functional networks. We studied this relationship between BOLD-fMRI and electroencephalography (EEG) signal of the human brain in detail using simultaneous fMRI and EEG in healthy awake human subjects at rest. Signals from EEG sensors were filtered into different frequency bands and reconstructed it in the three-dimensional source space. The correlation of the time courses of the two modalities were quantified on a voxel-by-voxel basis on full-brain level as well as separately for each resting state network, with different temporal shifts and EEG frequency bands. We found highly significant correlations between the BOLD-fMRI signal and simultaneously measured EEG, yet with varying time-lags for different frequency bands and different resting state networks. Additionally, we found significant negative correlations with a much longer delay in the fMRI BOLD signal. The positive correlations were mostly around 6-8 seconds delayed in the BOLD time course while the negative correlations were noticed with a BOLD delay of around 20 to 26 seconds. These positive and negative correlation patterns included the commonly reported alpha and gamma bands but also extend in other frequency bands giving characteristic profiles for different resting state networks. Our results confirm recent works that suggest that the relationship between the two modalities is rather brain region / network-specific than a global function and suggest that applying a global canonical HRF for electrophysiological data is probably insufficient to account for the different spatial and temporal dynamics of different brain networks. Moreover, our results suggest that the HRF also varies in different frequency bands giving way to further studies investigating cross-frequency coupling and its interplay with resting state networks.

## 1. Introduction

Functional magnetic resonance imaging (fMRI) based on the blood-oxygen-level-dependent effect (BOLD) is currently a cornerstone method in neuroscience. It is commonly applied to study the brain during rest and task; in healthy and diseased participants (Smith et al., 2009; Zhang & Raichle, 2010). Its biggest advantage is the spatial resolution unmatched by all non-invasive electrophysiological modalities. However, the BOLD signal is only an indirect measure of the underlying neuronal activity. It is presumed to be a result of a series of physiological events that follow neuronal activation including localized changes in cerebral blood flow, cerebral blood volume, and cerebral metabolic rate of oxygen and deoxyhemoglobin content (Buxton et al., 2004; Ogawa et al., 1990). Hence, several studies strived to characterize the neurophysiological correlates of the BOLD signal and its connectivity patterns using various electrophysiological techniques. However, our understanding is still rather incomplete despite these previous works. For instance, the canonical hemodynamic response function (HRF) is commonly used to account for the delay of the BOLD-signal regardless of brain region, frequency of electric activity and functional networks.

Pioneering animal studies measured BOLD and local field potential (LFP) power simultaneously in monkeys and cats and revealed consistently highly correlated LFP power with BOLD signal in the high gamma band mainly at 40 to 100 Hz (Logothetis et al., 2001; Niessing et al., 2005; Schölvinck et al., 2010; Shi et al., 2019; Shmuel & Leopold, 2008). This relationship was also found when correlating simultaneous BOLD and LFP signals in the human auditory cortex (Mukamel et al., 2005). Three of these animal studies also examined the time delay between the signal acquired from different modalities and demonstrated that the hemodynamic signal lagged the neural signal by 6 – 8 s (Logothetis et al., 2001; Schölvinck et al., 2010; Shmuel & Leopold, 2008). Additionally, Schölvinck et al. also reported a strong, positive correlation in lower frequencies (2–15 Hz) with a lag closer to zero.

Earlier human studies focused on correlating the BOLD signal to electroencephalography (EEG) using the occipital EEG electrodes and consistently reported negative correlations of the alpha band power between 8 – 12 Hz and BOLD signal (Goldman et al., 2002; Laufs, Kleinschmidt, et al., 2003; Laufs, Krakow, et al., 2003; Liu et al., 2012; Moosmann et al., 2003). Mantini et al. correlated EEG power variation from different frequency bands with that of fMRI resting state networks (Mantini et al., 2007) and found specific electrophysiological signatures for each network. Recently, another group studied the correlation between BOLD global signal and the global signal of simultaneously recorded EEG and reported both a high gamma correlation as well as negative correlations in the lower bands including alpha (Huang et al., 2018). These combined EEG-fMRI studies investigated correlations using electrode time courses and did not apply source-localization techniques. Therefore, a direct spatial correspondence between the sensor-space EEG results and fMRI is not possible. These mentioned studies also did not investigate the time-lag between the two modalities. Interestingly, Moosmann et al. studied simultaneous EEG-NIRS as well as simultaneous EEG-fMRI and reported a lag of about 8 s between simultaneous EEG-NIRS while such a lag was not reported in the simultaneous EEG-fMRI (Moosmann et al., 2003). Many of the human EEG studies applied a convolution of the EEG time courses with the HRF, in particular all studies reporting EEG negative correlation with fMRI used this approach. The use of HRF is a common procedure in neuroimaging to account for the delay between neural activation and its reflection in fMRI BOLD signal (Boynton et al., 1996). Classically it is assumed to be canonical, i.e. constant/identical in the whole brain. However, several experiments have reported variations of the HRF in different cortical regions, across subjects and in different measurement sessions (Aguirre et al., 1998; Handwerker et al., 2004; Miezin et al., 2000; Tewarie et al., 2016). Moreover, it is not clear if a canonical function is suitable to study neuronal/vascular coupling, i.e. the relation of EEG and BOLD signal. In summary, it is still unclear how neuroelectric activity and the BOLD response relate to each other.

In this work, we set to quantify and establish the link between spontaneous resting-state brain activity in simultaneously measured fMRI BOLD signal and source-reconstructed 256 channel high-density EEG (HD-EEG). The combination of this high number of electrodes and source-localization favors a more precise assessment of the dependencies of the two modalities in time and space and could aid in clarifying the still enigmatic BOLD-EEG coupling. We use data-driven analysis of the whole brain in different time-shift intervals from -30 to 30 seconds with minimal a priori constraints. We also evaluate this cross-modality coupling in different EEG frequency bands. Since functionally different brain regions differ in terms of their neuroanatomy and function, the relationship between EEG- and BOLD-signals might also vary depending on the brain regions. To test this hypothesis, in addition to a global characterization of the EEG-BOLD coupling, we assessed network-specific differences. We opted for studying resting state networks since each network connects different brain regions that are functionally related, and hence share similar dynamics of their time courses. We also evaluate the cross-modality coupling in different EEG frequency bands.

## 2. Materials and methods

### 2.1. Participants

We recruited 20 healthy participants without any history of neurological disorders. Five participants were excluded during the preprocessing due to artifacts (see fMRI and EEG data acquisition and preprocessing below), leaving 15 participants for further analysis (8 males, 7 females, mean age: 38.9 years, SD = 13.7 years, range 19-63 years). The study was approved by the Ethics committee of the Medical Faculty at the University of Tübingen and was conducted in accordance with the guidelines of the Declaration of Helsinki. All participants gave written consent before measurements.

### 2.2. Data acquisition

Magnetic resonance imaging data was acquired using a Siemens MAGNETOM Trio 3T scanner (Siemens AG, Erlangen, Germany) with a 12-channel array head coil for reception and the body coil for transmission. We acquired a sagittal T1-weighted volume with a 3D-MPRAGE sequence as high-resolution anatomical reference (TR 2.3 s, TI 1.1 s, TE 3.03 ms, FA 8°, voxel size 1 × 1 × 1 mm^3^); we also recorded a B0 field map for later correction of distortions in the functional images caused by magnetic field inhomogeneity (TR 2 s, TE 32 ms, FA 90°, 32 slices, voxel size 3 × 3 × 4 mm^3^). For the functional sequence, we acquired 180 gradient-echo planar T2^*^-weighted images covering the whole brain (TR 2 s, TE 32 ms, FA 90°, 32 slices, voxel size 3 × 3 × 4 mm^3^). The measurement duration was 10 minutes and participants were instructed to close their eyes and not to fall sleep.

Simultaneously with the fMRI measurement, we recorded a continuous high-density EEG signal using 256-channels EEG system (Electrical Geodesics, Inc., Eugene, OR, U.S.A.) and a sampling rate of 1000 Hz. Electrocardiogram (ECG) was measured simultaneously. The MR and EEG scanner clocks were synchronized, and the MR helium pump was turned off during the EEG resting-state measurement to reduce the noise induced by the pump in the EEG data (Nierhaus et al., 2013). This dataset will be referred to as <u>simultaneous-EEG</u> in this manuscript. We recorded an additional ten-minutes HD-EEG resting state measurement in an electrically shielded room outside the fMRI scanner. The second recording used the same 256-channel EEG system mentioned above. This dataset will be referred to as <u>non-simultaneous-EEG</u>. Both measurements were conducted in supine position.

### 2.3. Structural and functional MRI preprocessing

MRI processing was done in MATLAB (http://www.mathworks.com) using Statistical Parametric Mapping (SPM 12 [6470], Wellcome Trust Centre for Imaging Neuroscience; http://www.fil.ion.ucl.ac.uk/spm) as well as the FSL toolbox [5.0.9] (FMRIB, Oxford, UK; https://fsl.fmrib.ox.ac.uk/fsl). Using unified segmentation of SPM12, the structural T1-weighted images of all subjects were segmented into six tissue classes; grey matter, white matter, cerebrospinal fluid (CSF), skull, soft tissue outside the brain and finally air and other objects outside the head. The grey matter, white matter and CSF segmentations were joined to yield a segmented image of the intracranial volume (used as analysis space) and then spatially normalized and warped to MNI space using the DARTEL toolbox of SPM12. The flow fields of each participant’s anatomical transformation to the DARTEL template in MNI space were later used for warping the functional data. Tissue masks were also generated by binarizing the segmented tissues at a threshold of 0.5.

fMRI functional time series were first slice-time corrected using the first slice (slice-time = 0 ms) as reference to correspond with the TR trigger sent to the EEG and conserve temporal comparability of the two methods. Head motion correction was then performed using FSL MCFLIRT (Jenkinson et al., 2002). Additionally, a framewise displacement (FD) (Power et al., 2014) threshold was set to 0.5 mm and volumes exceeding this threshold were discarded. Figure S1 shows the fMRI volumes framewise displacement for each subject along with the scrubbed above-threshold frames. The first ten volumes of fMRI data were also removed to avoid T1 effects. Volumes that were either scrubbed in fMRI or corresponded to EEG-artifacts from both modalities were discarded (see details on EEG processing below). This was done to preserve the temporal comparability of the fMRI and EEG data.

The voxel displacement maps (VDM) were then calculated using the acquired fieldmap images and the Fieldmap Toolbox [version 2.1] integrated in SPM. The time series were then distortion-corrected, realigned, co-registered to the T1-weighted anatomical reference image (normalized mutual information cost function) and finally normalized to MNI space using the DARTEL flow fields (compare above). The normalized data were smoothed with an isotropic Gaussian kernel (5 mm full width at half maximum). Individual brain masks were also normalized to MNI space using the DARTEL flow fields. Afterwards, a high-pass filter of 0.1 Hz was applied to the data. Finally, the normalized functional datasets were masked with the individual brain mask and a global brain mask (generated by averaging all normalized individual masks and then binarizing at the threshold of 0.8). This procedure ensures that only brain voxels that are consistently present in at least 80% of all subjects were considered in the further analysis.

### 2.4. Resting States Networks masks extraction

To extract data-driven fMRI resting state networks we used the CONN connectivity toolbox [version 18b] (http://www.nitrc.org/projects/conn) (Whitfield-Gabrieli & Nieto-Castanon, 2012) on preprocessed fMRI time courses to extract 40 ICA components using the FastICA algorithm. The resulting components masks were spatially matched with seven reference resting state networks obtained from the Yeo et al. (Yeo et al., 2011) resting state atlas namely visual, somatomotor, dorsal attention, ventral attention, limbic, frontoparietal and default mode networks. This spatial matching was performed with the FSLCC function in FSL with a correlation threshold of 0.2. Components surpassing the spatial correlation threshold are considered part of the network and were spatially merged into one mask for that network. See figure S2 for data-driven topographical maps of the extracted fMRI networks.

### 2.5. EEG preprocessing

MR gradient artifacts due to static and dynamic magnetic fields were removed from the simultaneously measured EEG data using average artifact subtraction (AAS) method (Allen et al., 2000). Cardioballistic artifacts were detected and removed by the Pulse Detection Tool implemented in Net Station 5.2 software (Iannotti et al., 2015). The data was then downsampled to 250 Hz, demeaned and band-pass-filtered at 0.2-100 Hz. Power-line artifacts were removed by a bandstop filter of 49.5 to 50.5 Hz. The data was then segmented into epochs of 2 seconds where the fMRI TR trigger corresponded to the center sample (epochs ±1 s around fMRI trigger). The data was visually inspected to identify and discard noisy electrodes and epochs. Note that discarded artifacts in simultaneous-EEG were also removed from fMRI (compare to above). Preprocessing and analysis of EEG data was performed using the Fieldtrip toolbox (Oostenveld et al., 2011) running in MATLAB (version 9.0 [R2016a] Mathworks Inc.). See Table S1 for a comprehensive list of the number of channels and epochs removed from each subject’s data. In all modalities, at least 86% of the data and 80% of the electrodes were admitted to further analysis.

### 2.6. Correlating EEG Alpha power with BOLD signal

As a sanity check for our data and pipeline, we aimed at reproducing the analysis and findings employed by Laufs et al. in 2003 studying the relationship between fMRI and EEG Alpha power derived from occipital, central and frontal electrodes. We followed the same preprocessing steps as Laufs et al. 2003 but implemented it in Fieldtrip. We took the arithmetic mean of the two occipital electrodes corresponding to O1 and O2. The same was done for parietal electrodes C3/C4 and frontal ones F3/F4 for control purposes. The powers of EEG timecourses were then limited to their mean plus or minus 3 SD to account for brief motion or muscle artifacts. We performed Fast Fourier transformation using a hanning window on two second epochs. The spectral power was then demeaned and averaged across the alpha band frequency bins. Using this filtered Alpha signal per 2 seconds epochs (matching the TRs of the BOLD), we generated an additional version of the data by convoluting it with the HRF function. Each dataset (with and without HRF-convolution) was then correlated with the fMRI BOLD time-courses at each voxel obtaining a correlation brain map. The correlation was tested among the three types of electrodes using a Kruskal-Wallis-Test. Significant voxels were separated into negative and positive ones based on the observed median correlation at that voxel. We then extracted clusters with minimum 9 neighboring voxels using the MATLAB function “bwlabeln” and three-dimensional 18-connected neighborhood (separate clusters for positive and negative correlations). The median correlation for significant voxels was summed across clustered significant voxels in each of the 7 resting state networks and normalized by the number of voxels in that network. Figures S3 and S4 show the normalized summed correlations observed in each network for both positive and negative correlations.

### 2.7. EEG source reconstruction

The preprocessed artifact-corrected EEG data was filtered into seven different frequency bands: delta (1-4 Hz), theta (4 – 8 Hz), alpha (8 – 12 Hz), beta1 (12 – 20 Hz), beta2 (20 – 30 Hz), gamma1 (32 – 48 Hz), and gamma2 (52 – 68 Hz). Filtered data in each frequency band was projected onto a regular 5-mm grid spanning the entire brain using linearly constrained minimum variance (LCMV) scalar beamformer (Veen et al., 1997). For each grid position, the leadfield was calculated using a boundary element model (BEM) constructed from the participant’s structural MRI where the participant’s scalp, skull and brain surfaces were modeled. The first two boundaries, namely the scalp and the skull, were derived from corresponding SPM segmentations (c5 and c4 respectively). The brain surface was derived from the combination of the grey matter, white matter and CSF segmentations. We also calculated a covariance matrix for each frequency band. Using both the leadfield and the covariance matrix, a spatial filter was calculated for each of the studied frequency bands and used to project the time courses by the EEG sensors into source-space. For each grid-position, the amplitude envelope was computed as the positive magnitude of the Hilbert transformation of the signal (Hilbert envelope). The Hilbert envelope was low-pass filtered to 0.25 Hz and then downsampled to 0.5 Hz. To ensure that the fMRI and MR-EEG time courses were temporally comparable, we selected the EEG samples that corresponded temporally to the registered TR-pulse triggers. The source-space EEG signal time course for each frequency band was exported in NIFTI format, up-sampled to a voxel size of 3 mm. It underwent the same spatial processing as the fMRI data, i.e. the volumes were warped into MNI space (via DARTEL flow fields), masked with individual brain masks and finally masked with a global brain mask. Non-simultaneous-EEG data acquired outside the scanner was preprocessed in the same pipeline as the simultaneous-EEG data except for the removal of MR gradient and cardioballistic artifacts.

### 2.8. Correlation of time courses

To study the relationship between fMRI and EEG signals in different frequency bands, we studied the statistical dependency between the fMRI and different frequency specific EEG time courses for each voxel. We used Pearson’s correlation coefficient between time courses of both modalities at each voxel. Furthermore, we investigated these correlations at different temporal shifts between +30 to -30 seconds (cross-correlation). By shifting the fMRI backwards, we studied the assumption that neural signals are reflected in fMRI signal later than in EEG signal. Shifting the fMRI time course forward investigates the contrary. We studied both backward and forward lags of fMRI time courses from 1 to 15 samples, which represents 30 seconds in both directions in steps of two seconds (TR = 2 seconds). We then calculated Pearson’s correlation coefficient of the time course in each of the fMRI shifts with each of the EEG time courses (simultaneous- and non-simultaneous EEG, filtered in 7 frequency bands each). This procedure yielded a correlation matrix for each voxel per subject, frequency band, time-lag, and EEG dataset (simultaneous- and non-simultaneous-EEG). See figure 1 for a schematic depiction of the analysis pipeline.

**Figure 1:**
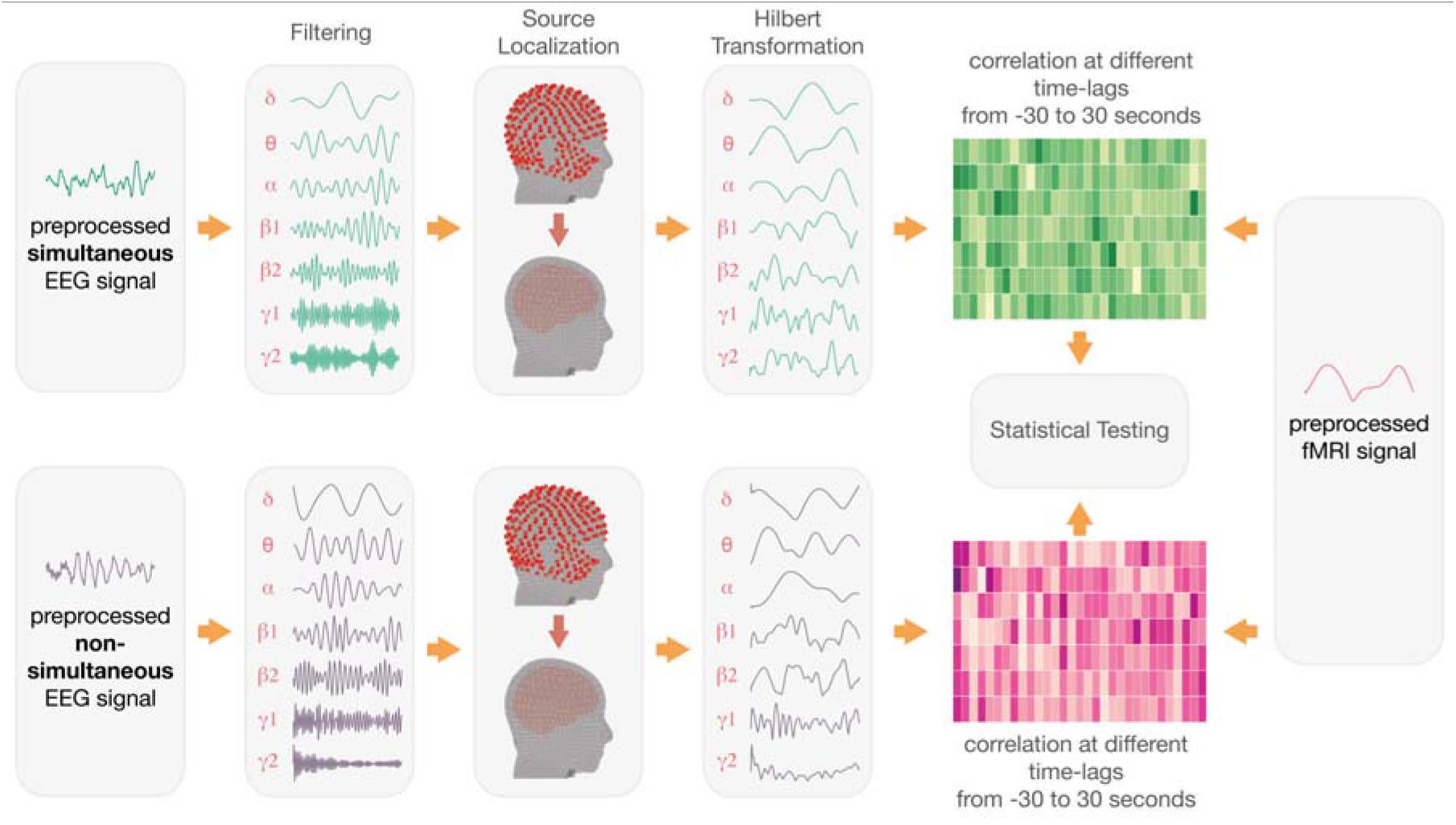
Outline of the steps used to obtain the correlations from preprocessed fMRI and EEG time courses. See figure S5 for the statistical testing pipeline.

### 2.9. Statistical Testing and Network Analysis

Since we assumed the absence of any meaningful temporal dependency between fMRI and non-simultaneous EEG, we considered their correlations as a control condition. This approach ensured that the general characteristics of the EEG signal (autocorrelations, microstates, etc.) should be similar in both conditions, since the EEG was measured in the same subject with the same system. Thus, our null hypothesis is that simultaneous EEG to fMRI correlations do not differ from non-simultaneous EEG to fMRI. A significantly higher simultaneous EEG to fMRI correlation would be perceived as a positive correlation between the two modalities while a significant lower correlation would be considered as a negative correlation (anticorrelation).

The statistical analysis was done in several steps, as follows:

Step 1: First we performed a first-level between groups non-parametric Wilcoxon signed-rank test on the correlation values of simultaneous EEG:fMRI and non-simultaneous EEG:fMRI at each voxel and obtained a z-score.

Step 2: The sum of positive z-scores of significant voxels (alpha = 0.05) divided by the sum of tissue probability weights. This was done to account for outlier values on the edge of the tissue / network boundaries. The weighted value was then noted per frequency band and time-lag yielding a 7×31 matrix (7 frequency bands and 31 time-lags from -30 to 30 s in steps of 2 s). Similarly, another matrix was generated by summing only negative z-score values to capture negative correlations.

Step 3: To test patterns of significant difference between the fMRI correlation with simultaneous vs. non-simultaneous EEG, we used a permutation-based approach to generate control-matrices by repeating the previous steps after randomly re-assigning the correlation values to one of the two conditions (simultaneous vs. non-simultaneous). We performed 10,000 permutations and in each we separately calculated the weighted sum of positive z-scores and the weighted sum of negative z-scores. For each frequency / time-lag combination, we set an arbitrary threshold level for a z-score sum at the 90^th^ percentile of the 10,000 values.

Step 4: Using these thresholds obtained from step 3 (7×31 matrix of 7 frequency bands and 31 time-lags) we binarized the observed matrices as well as the permutation matrices to prepare for clustering analysis.

Step 5: We assumed that a biologically meaningful relationship between fMRI and EEG would not be limited to one specific frequency band/time-lag combination but would rather be present in neighboring frequencies and/or time-lag combinations as well. Therefore, we used a cluster size of minimal 3 neighboring matrix positions as a threshold to extract clusters in each binarized matrix. The neighborhood was based on the “two-dimensional four-connected” principle using the “bwlabeln” function in MATLAB. For each observed cluster, we calculated the sum of the statistical values present of all points in the cluster (cluster sum score).

Step 6: From each permutation, we used the maximal cluster sum score and constructed a random distribution for the null-hypothesis. The observed cluster sum score of the un-permuted experiment was then tested against this null-curve of 10,000 permutations to determine the error probability that the actually found cluster value was generated by chance. The frequency of equal or higher values was reported as the probability (p) value of that cluster. See figure S5 for an illustration of the statistical approach employed.

To assess the globality of the relationship between the two modalities, we performed the previous steps on different sets of voxels (regions of interest = ROI); namely all grey matter, white matter and CSF voxels as well as on the voxels belonging to each resting state network separately (0.5 ROI mask threshold). For each of those ROIs, we generated the positive and negative matrices of sum z. The alpha level for the cluster statistics (step 6) was set at p=0.0025 (0.5/20 tests) to correct for multiple testing at 7 networks and 3 tissue types (grey matter, white matter and CSF) in 2 different matrix types (positive and negative).

### 2.10. Testing network pattern differences

To test whether the extensions of the cluster patterns were different between networks, we determined the centroid of each cluster and calculated its distance to all points that form the cluster polygon. For each pair of clusters, we pooled the distances from both clusters together in order to get a probability distribution for distances in this cluster pair. We tested the distance between the two centroids against the distance distribution by determining the probability of the inter-centroid distance being within the range of either cluster. We used an alpha level of p = 0.05 to determine significantly different/distant clusters. We performed this step for each pair of tested networks positive correlation matrices as well as between pairs of negative correlation matrices.

### 2.11. Data and code availability

All data and code used in this study will be available upon reasonable request to the authors.

## 3. Results

We first studied the correlation between the fMRI BOLD and the EEG Alpha power from occipital, central and frontal electrodes. This was done to reproduce previous findings using our pipeline and as a sanity check for the data quality particularly after rejection of various MR-related artifacts. Comparing correlation profiles from different resting state networks with BOLD time courses (without HRF convolution), we found the highest positive correlation with the EEG alpha power between the occipital electrodes and frontoparietal network, followed by default mode-networks (figure S4), while the strongest negative correlation was found in the same networks respectively but with the frontal electrodes. When the BOLD time courses were convoluted with the HRF, we found the strongest positive correlation between the visual network and the frontal electrodes while the strongest negative correlations were between the visual network and the occipital electrodes (figure S3).

Next, we studied the global relationship between the fMRI and simultaneous-EEG in the gray matter, which is the main finding in this work. We tested this correlation in each frequency band in time-lags between -30 and +30 seconds (figure 2) using the source-localized EEG activity. On the one hand, we found a cross-spectral pattern of positive correlations spanning all the studied frequency bands (p<0.0001). The maximal positive correlation was found in the high gamma band with around 6 – 8 seconds lag of the BOLD time courses in relation to the EEG signal. Interestingly, correlations in the lower frequencies had their maxima at around 2 seconds. On the other hand, we found two clusters of negative correlations; one in low frequencies with a maximal negativity in the delta band with a lag of about 20 seconds (p<0.001) and a second cluster extending towards the higher frequencies with a maximal negativity in the lower gamma band and a time-lag of around 24 seconds (p<0.0001).

**Figure 2:**
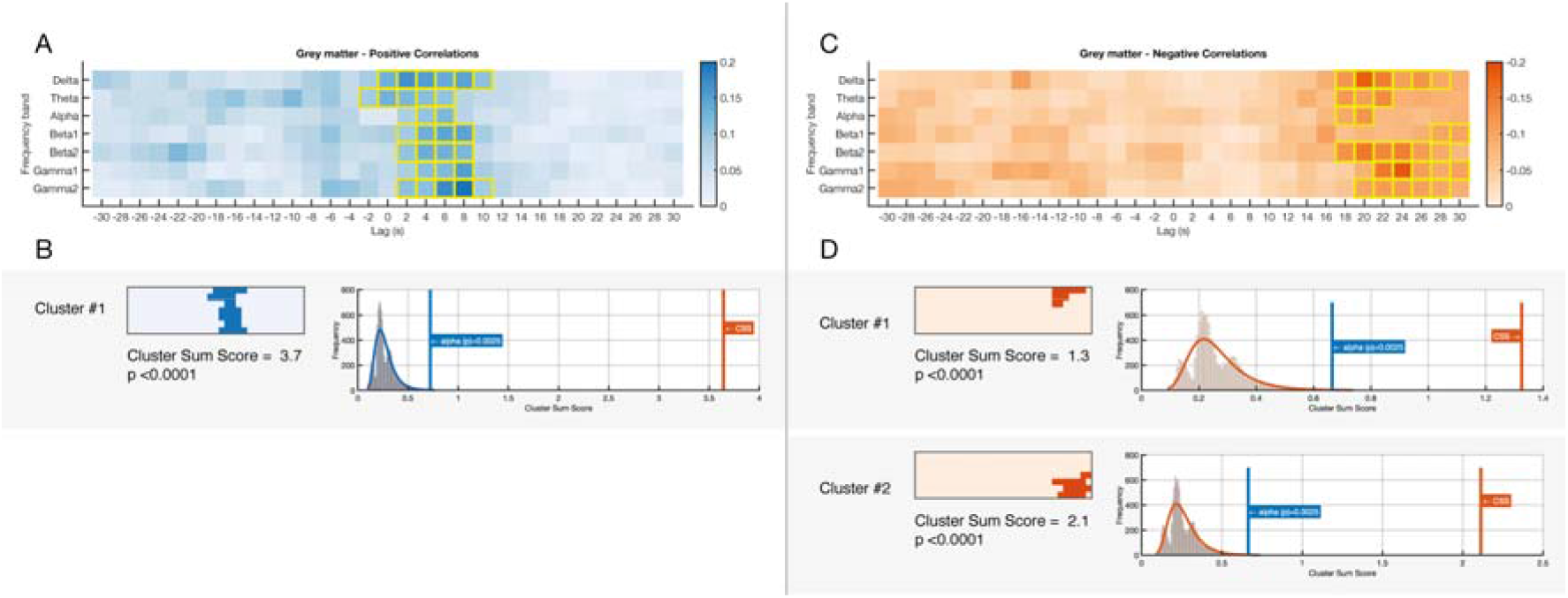
EEG-fMRI significant correlations across frequencies and time-lags in grey matter. (A) Shows a matrix of positive sum-z scores derived from significant positive correlations in the first-level statistics. Significant clusters are outlined with yellow strokes. (B) Significant clusters derived from positive correlations in A along with their *cluster sum score* compared to the random distribution derived from 10,000 permutations where the two conditions were randomly shuffled. C and D: Similar sections to A and B but showing results derived from significant negative correlations.

We also calculated the previous matrices separately for the white matter and CSF and found weaker but similar effects. However, the white matter positive correlations matrix was only significant in the higher frequencies (p<0.0001) and the CSF correlations were maximum in the lower frequencies (p<0.0001) (figures S6 and S7).

To test whether the effect in the gray matter is stereotypical across the whole brain or if there are regional/network differences, we tested the correlation patterns for gray matter separately for the 7 resting state networks. We generated matrices for both positive and negative weighted (sum z) of the correlations for each network separately. We also performed the permutation-based statistical assessment separately on each network (Bonferroni corrected for all tests). Patterns of positive significant clusters were found in all networks except for the dorsal attention network while significant negative clusters were found in four networks namely the visual, somatomotor, dorsal attention and limbic networks (figure 3 for the networks matrices and the figures S8 to S14 for clusters random distribution plots). Consistently, positive correlations preceded the negative correlations in all networks. We did not find negative correlation clusters earlier than 10 seconds lag and the positive correlations were lagged with maximum 16 seconds as in the visual network.

**Figure 3:**
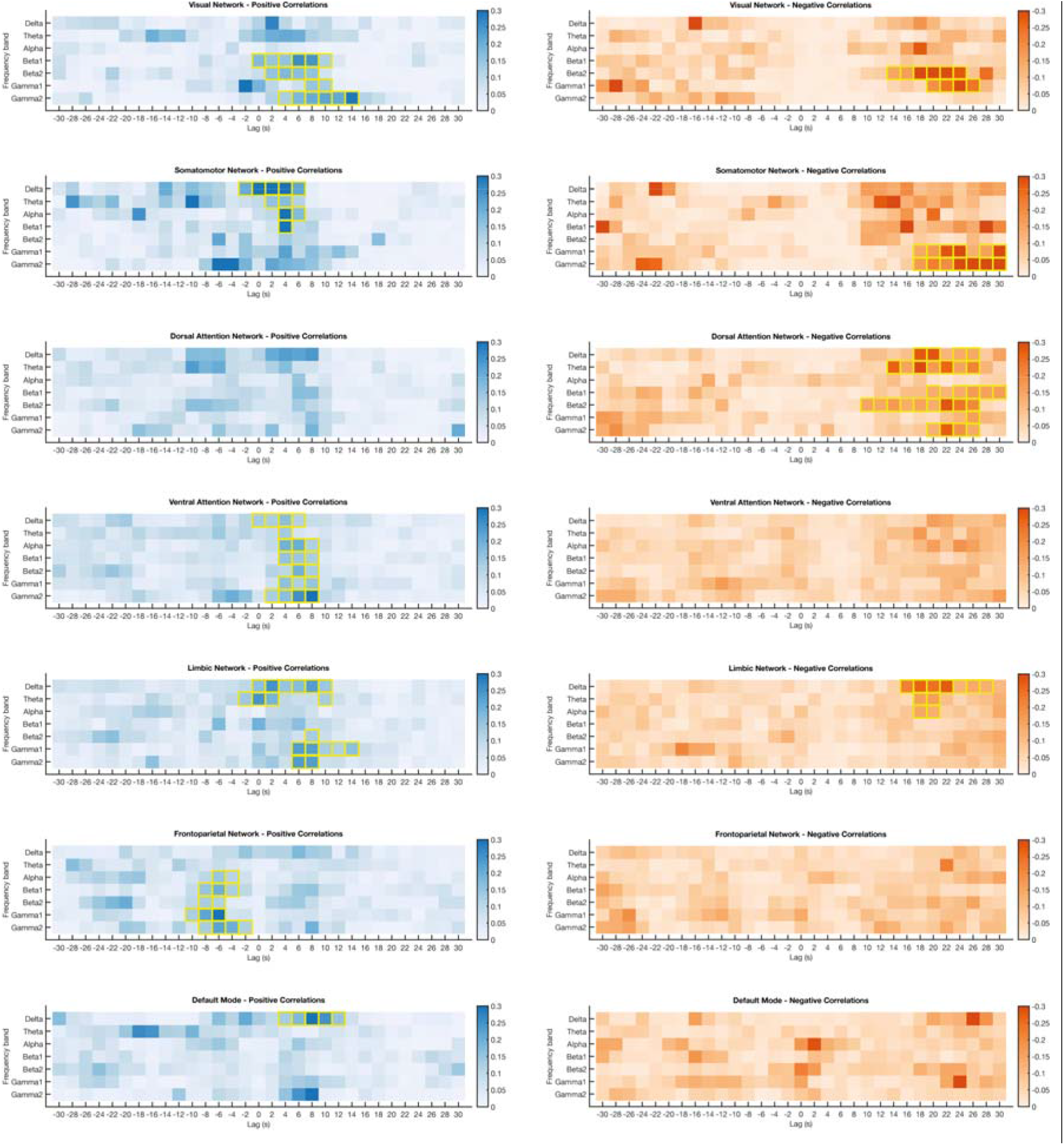
fMRI:simultaneous-EEG correlations across frequencies and time-lags in different resting state networks. Significant clusters are outlined with yellow strokes.

The correlation patterns were not uniform across all resting-state networks. We found significant differences in the temporal/frequency matrix in 9 positive comparisons out of 21, mainly between the frontoparietal network and other networks. In the negative correlation patterns, we found 2 significant pairwise correlations out of 10 comparisons. These differences were between the limbic and dorsal attention network on one hand and the somatomotor network on the other hand. (see figure 4 for a schematic plot of the similarities indices as well as estimated significance levels. See figures S15 and S16 for more detailed depiction of the clusters relationships).

**Figure 4:**
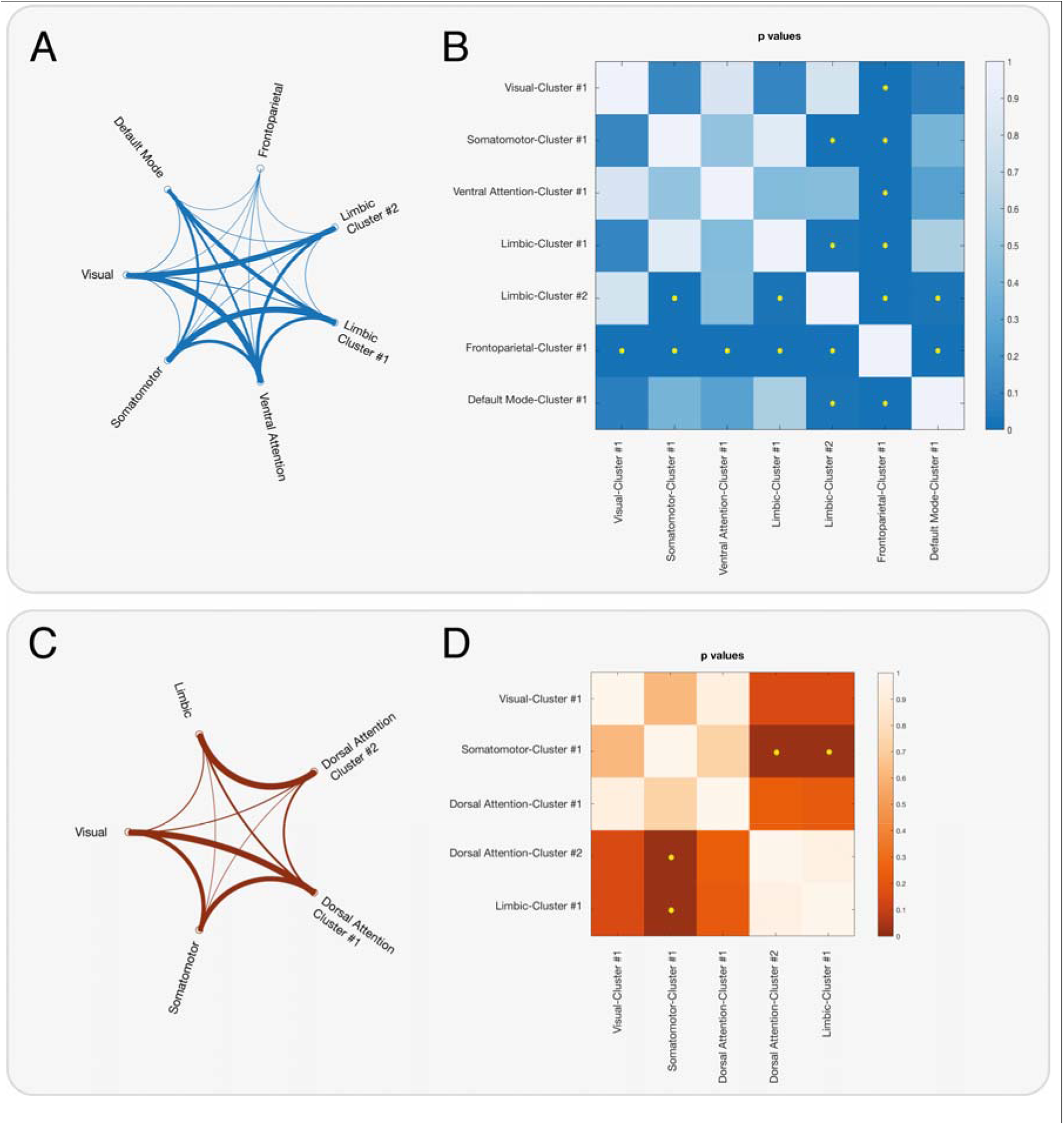
Visualization of the correlation similarities between networks temporal-frequency band clusters. A: Pairwise network similarity (1-*p*-*value*) based on Euclidean distance between their respective significant positive clusters in the temporal-frequency band profile. Connection thickness is inversely proportional with the probability that the correlation pattern in the connected cluster is different. B: Matrix showing the p-values of pairwise comparisons of the clusters. Significantly different clusters comparisons are marked with yellow stars (alpha = 0.05). C and D: As in A and B respectively but using significant negative clusters. Note that only significant clusters were included in this analysis, hence the difference in the number of comparisons between positive and negative correlations.

## 3. Discussion

We investigated the spatiotemporal coupling of the fMRI-BOLD and EEG signal in a data-driven approach. Our results demonstrate a highly significant correlation between the fMRI signal and simultaneously measured EEG that varies in frequency bands and time-lags as well as between resting state networks. Globally, a significant positive correlation was found with fMRI lagging behind the EEG signal. This lag, however, varied in different EEG frequency bands with lower EEG frequencies having shorter fMRI to EEG lags (2 to 4 seconds) and higher EEG frequencies having longer lags (6 – 8 seconds). The positive correlation in higher EEG frequency bands are consistent with findings reported from various studies investigating BOLD and simultaneous electrophysiological measurements (Lachaux et al., 2007; Scheeringa et al., 2011; Schölvinck et al., 2010; Shmuel & Leopold, 2008). On the other hand, the lower EEG frequency band correlations are similar to those previously observed by Schölvnick et al. while studying BOLD and cortical LFP in monkeys (Schölvinck et al., 2010). This frequency-shift combination seems to vary as well in different resting state networks suggesting that the electrophysiological correspondence of fMRI BOLD signals is a rather multifaceted than a stable global relationship.

Neuronal activities are reflected almost instantaneously in electrophysiological recordings (EEG in our case). Conversely, the reflection of these activities in BOLD oscillations has a longer temporal delay in the range of several seconds. The canonical hemodynamic response function (HRF) is, therefore, commonly used in general linear models to relate BOLD responses with experimental stimuli or EEG derived metrics (like power). However, several studies have suggested that the HRF and consequently the temporal delay in fMRI vary across brain regions and in different individuals (Aguirre et al., 1998; Handwerker et al., 2004; Miezin et al., 2000; Taylor et al., 2018). Our results confirm that the relationship also varies among different resting state networks. Moreover, we hypothesized this relationship to vary in different EEG frequency bands. In fact, some networks exhibited maximal correlation in low frequency bands like delta and theta while other networks tended to correlate more in higher frequencies like beta and gamma. This is generally in line with various studies showing that hemodynamic response following neural activities varies in different networks and cortical locations (Aguirre et al., 1998; Hipp & Siegel, 2015; Mantini et al., 2007). Since the function of each network appears to be formed by a collective of cross-spectral activation patterns (Mantini et al., 2007), it remains difficult to attribute the different time-lags to either functional network, the dominant frequency band or the interplay of these two aspects.

Given variable temporal delays in different EEG frequency bands and different networks, our results and the available literature challenge the notion that a simple and universal model of a canonical function for EEG-fMRI/BOLD coupling is valid for all cortical regions and all spectral frequencies. Since employing a non-specific HRF model could lead to dramatic changes in the resulting BOLD activation and its temporal precision, there is an urgent need to further elucidate this relationship and possibly update current standard imaging analysis methods. Regional, or frequency-band specific data-driven HRF generation could offer a potential solution at least when studying EEG-fMRI coupling.

In our results, we also observed patterns of negative correlation between the two modalities that temporally followed the positive correlations and lagged around 20 seconds in general. One possibility is that this negative correlation reflects the undershoot seen in the HRF which have been related to delayed vascular compliance and sustained increases in the metabolic rate of oxygen (van Zijl et al., 2012).

Unlike previous simultaneous EEG-fMRI studies (Goldman et al., 2002; Huang et al., 2018; Laufs, Kleinschmidt, et al., 2003; Liu et al., 2012; Moosmann et al., 2003), we did not observe a strong negative correlation between the modalities in the alpha band especially in the occipital lobe using our main voxel-to-voxel analysis. We only found a relatively weak negative correlation for the alpha in the default-mode network sub-analysis around 2-4 seconds that did not survive the cluster-level correction (compare figure 3) and was not evident in the global analysis. However, we also observed a negative correlation of the BOLD fMRI with the occipital alpha band EEG derived from the surface electrodes after convolution with the HRF. Since we did not use a canonical HRF to convolve the EEG time courses in our main voxel-to-voxel analysis, we could postulate that Applying a canonical HRF could possibly alter the temporal characteristics of the EEG signal and produce imprecise correlations with the corresponding BOLD signal. Such signal convolution was also not implemented in the typical animal studies (Logothetis et al., 2001; Schölvinck et al., 2010; Shmuel & Leopold, 2008).

Many animal and human studies employing various electrophysiology techniques demonstrated that the gamma band presented the most common correlation with the BOLD signal in task and rest. This was done either by correlating time courses of fMRI and electrophysiological measures or by studying within modality connectivity and comparing the between modality similarities. While electrocorticography (ECoG) studies reported similarity in both low and high frequencies including gamma (Hacker et al., 2017; He et al., 2008; Keller et al., 2013), non-invasive studies reported mainly effects around the alpha and beta bands using EEG and magnetoencephalography (MEG) (Brookes et al., 2011; Deligianni et al., 2014; Hipp & Siegel, 2015; Tewarie et al., 2016). It is unclear whether this can be attributed to the physical limits of the method to capture the activity of specific neural generators, i.e. as is the case with MEG, which is insensitive to radial currents (Hacker et al., 2017; Mosher et al., 1992). In our study, we could confirm the gamma band correlation but also the presence of significant correlations in lower EEG frequency bands. Compared to the other studies, we used source-space localized EEG signals from 256 electrodes exceeding the maximum number of 64 electrodes in previous studies, which were also only investigated in the sensor space.

Various decisions in the data processing pipeline have to be done. For example whether to regress out the global signal (time series of averaged signal intensity across all brain voxels) from fMRI timeseries prior to correlating it with the EEG data remains unclear given the controversy surrounding global signal regression in the field (Fox et al., 2009; Murphy et al., 2009; Murphy & Fox, 2017; Power et al., 2017; Saad et al., 2012). In this work, we decided to preserve the global signal motivated by several recent studies that deduced its accountability for some electrophysiological correlates (Huang et al., 2018; Schölvinck et al., 2010; Wong et al., 2013, 2016). Nevertheless, a clear understanding of the contribution of the global signal to the electrophysiological signature of the BOLD response needs to be elucidated in further studies.

Interestingly, we have also observed patterns of lagged correlation in the white matter and CSF. Correlations between BOLD fluctuations in the CSF and white matter were also reported in previous studies (Fultz et al., 2019; Schölvinck et al., 2010). Recent works by Li et al. investigated the HRF in the white matter in response to stimulus and reported that the presence of white-matter specific HRF (Li et al., 2019). Fultz et al. studied electrophysiological, hemodynamic and CSF oscillations during sleep and found coherent dynamics of the three signals, which was also negatively related to the dynamics in the grey matter (Fultz et al., 2019). From our data, we cannot infer the nature of the observed effect in the white matter and CSF. It could possibly be attributed to the limitation of the EEG-source localization spatial-resolution.

## Limitations

A main obstacle in simultaneous EEG-fMRI experiments is the reduction of the MR artifact contaminating the EEG signal. We tackled this issue by turning off the helium pump and reducing the movement of the subject to the minimum during the acquisition as well as using state-of-the-art post-acquisition data preprocessing and artifact rejection methods. Since our results demonstrate correlation values at various, non-zero time-lags, it is unlikely that the observed correlations are largely due to residual gradient or other noise (movement, pulse, etc.), which would be expected to have a zero-lag between the two signals.

Another limitation in multi-modal studies is the combined limitation of each modality. For example, we had to downsample the EEG time courses to that of the fMRI discarding much of the high temporal information contained in the EEG. Emerging new fMRI sequences in fast-band fMRI (Chen et al., 2020; Sahib et al., 2016) would aid in the multimodal imaging field to improve the temporal resolution of fMRI and allow more fine-grained assessment of the dynamics of EEG-fMRI coupling. On the other hand, due to the volume conduction and the relatively low resolution of EEG source localization, the spatial precision of the correlation of the two modalities is generally limited. However, we can assume that source-reconstructed 256-channel EEG has reasonable spatial precision. In one of our recent studies, it was comparable to fMRI if an individual head-model was used (Klamer et al., 2015). Also studying the link between EEG and other imaging probes of brain function and metabolic markers ([^18^F]FDG-PET for glucose metabolism, [15O]H_2_O-PET for perfusion) may be of value.

## Conclusions

In this study, we used simultaneous fMRI and high-density EEG to investigate the relationship of neuronal and vascular/BOLD signal in a data-driven approach with minimal a-priori assumptions. We measured another non-simultaneous EEG dataset from the same subjects as a control condition and used source reconstruction of EEG signals for a better estimation of the spatial relationship between the two signals. We showed that neuronal/vascular coupling has a distinct temporal profile for different EEG frequency bands and for different resting state networks. These results show that the use of canonical “hemodynamic response functions” is not adequate for EEG-fMRI coupling and that network and frequency band specific effects need to be considered. Based on this work, we recommend using data-driven HRF for different brain regions or networks in different frequency bands.

The present findings provide a basis for further studies approaching the neural correlates of BOLD signal as well as studies seeking a deeper understanding of the mechanisms driving cross-frequency and cross-modality coupling.

## Acknowledgements

We thank all volunteer controls for participating in this study. We also thank Dr. Markus Siegel, University of Tübingen, for the fruitful discussions. The authors acknowledge support by the state of Baden-Württemberg through bwHPC. AE was supported by Center of Integrative Neuroscience, Tübingen (CIN) grant 2014-02. NKF was supported by DFG grant FO750/7-1. HFW was supported by DFG grant WE 5795/2-1, JSPS grant PE15015 and IZKF Juniorgrant (2209-0-0)

## Author Contributions

Conceptualization, A.E., C.P., and N.K.F.; Methodology, A.E., C.P. and N.K.F.; Software, A.E.; Investigation, A.E., S.K. and H.W.; Writing – Original Draft, A.E., C.P. and N.K.F.; Writing – Review & Editing, A.E., S.K, D.v.d.V, H.L., C.F., C.P. and N.K.F.; Funding Acquisition, N.K.F., C.F. and H.W.; Supervision, N.K.F, C.P., and H.L.

