## Supplementary Material for "Linking neuronal and hemodynamic network signatures in the resting human brain"

### Supporting Information

| Subject | First 10 Volumes from fMRI | fMRI volumes scrubbed | EEG epochs with artifacts | No. of discarded electrodes in EEG | No. of remaining volumes in both modalities |
| --- | --- | --- | --- | --- | --- |
| 1 | 10 | 2 | 5 | 45 | 284 |
| 2 | 10 | 0 | 6 | 31 | 284 |
| 3 | 10 | 25 | 30 | 36 | 258 |
| 4 | 10 | 0 | 3 | 48 | 287 |
| 5 | 10 | 17 | 26 | 24 | 264 |
| 6 | 10 | 0 | 1 | 54 | 289 |
| 7 | 10 | 4 | 12 | 29 | 278 |
| 8 | 10 | 18 | 20 | 20 | 267 |
| 9 | 10 | 10 | 18 | 8 | 272 |
| 10 | 10 | 1 | 5 | 52 | 285 |
| 11 | 10 | 3 | 5 | 9 | 284 |
| 12 | 10 | 3 | 8 | 27 | 282 |
| 13 | 10 | 2 | 8 | 44 | 282 |
| 14 | 10 | 0 | 5 | 18 | 285 |
| 15 | 10 | 29 | 32 | 42 | 258 |
| Table S1: Number of removed volumes/epochs from fMRI and EEG data respectively as well as discarded electrodes in EEG per subject. | | | | | |

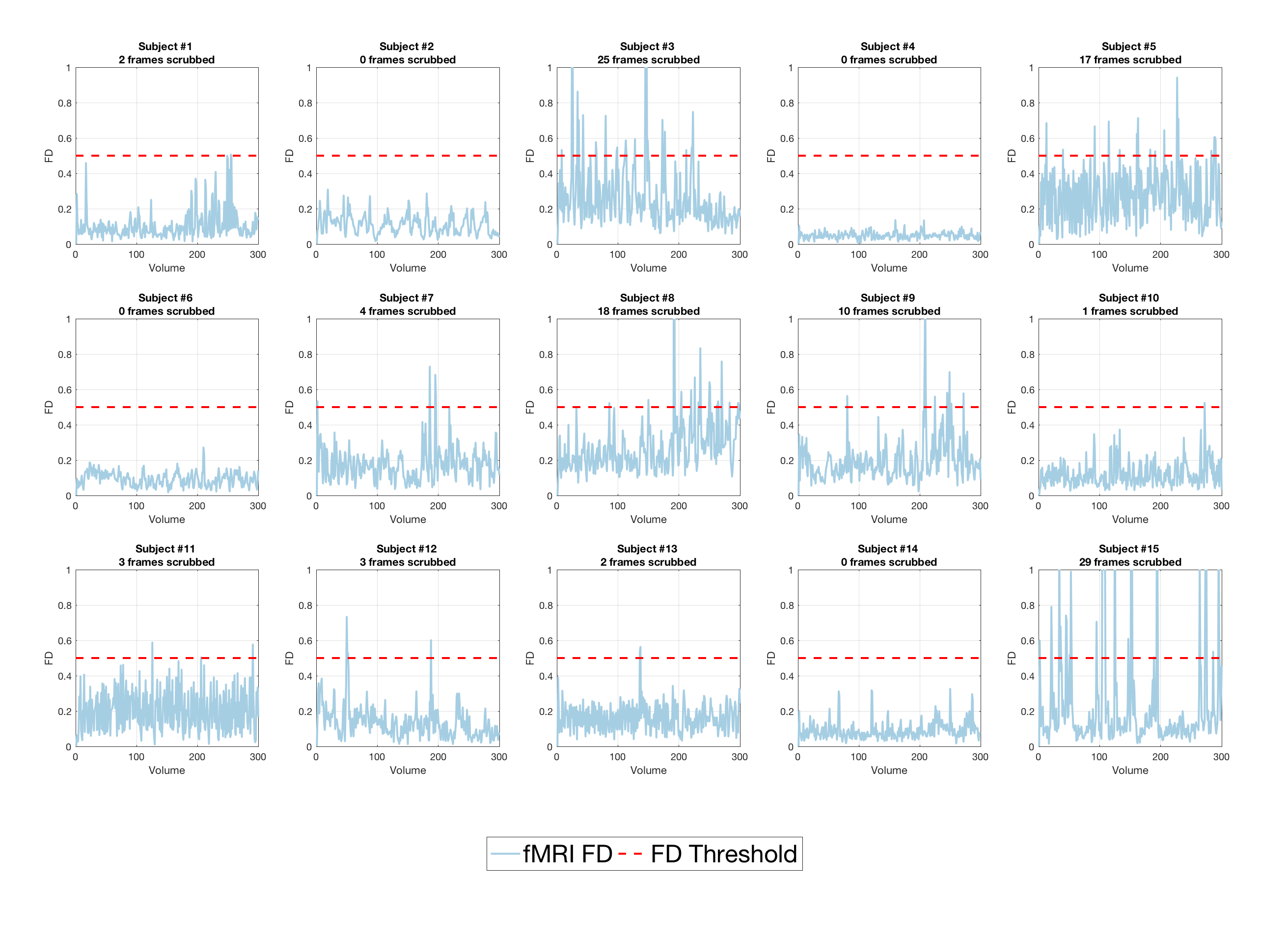

**Figure S1:** fMRI volumes framewise-displacements for each subject along with the scrubbed above-threshold frames. FD-Threshold = 0.5. (FD: Framewise Displacement)

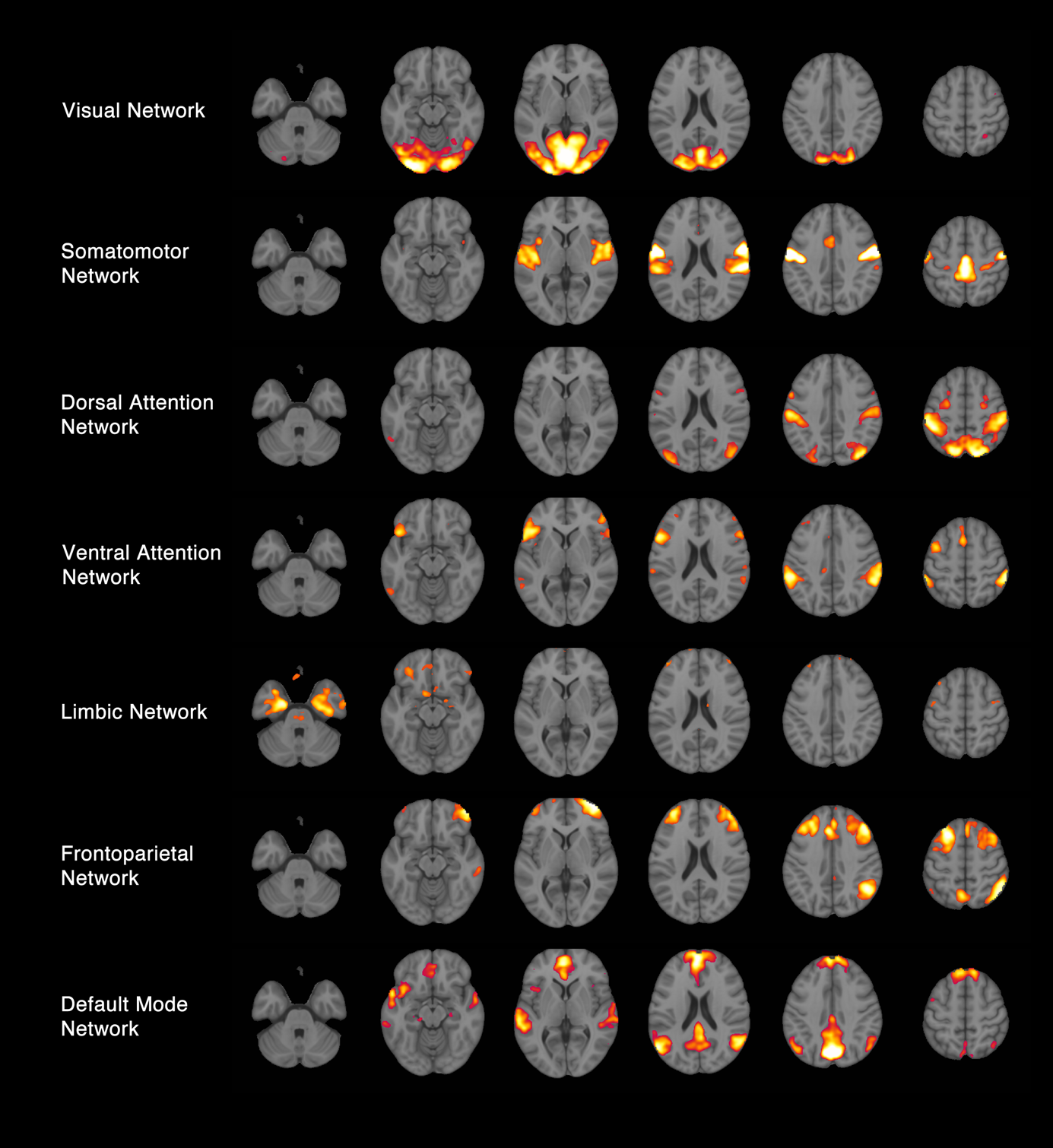

**Figure S2:** Data-driven resting state networks matching reference networks from (Yeo et al., 2011).

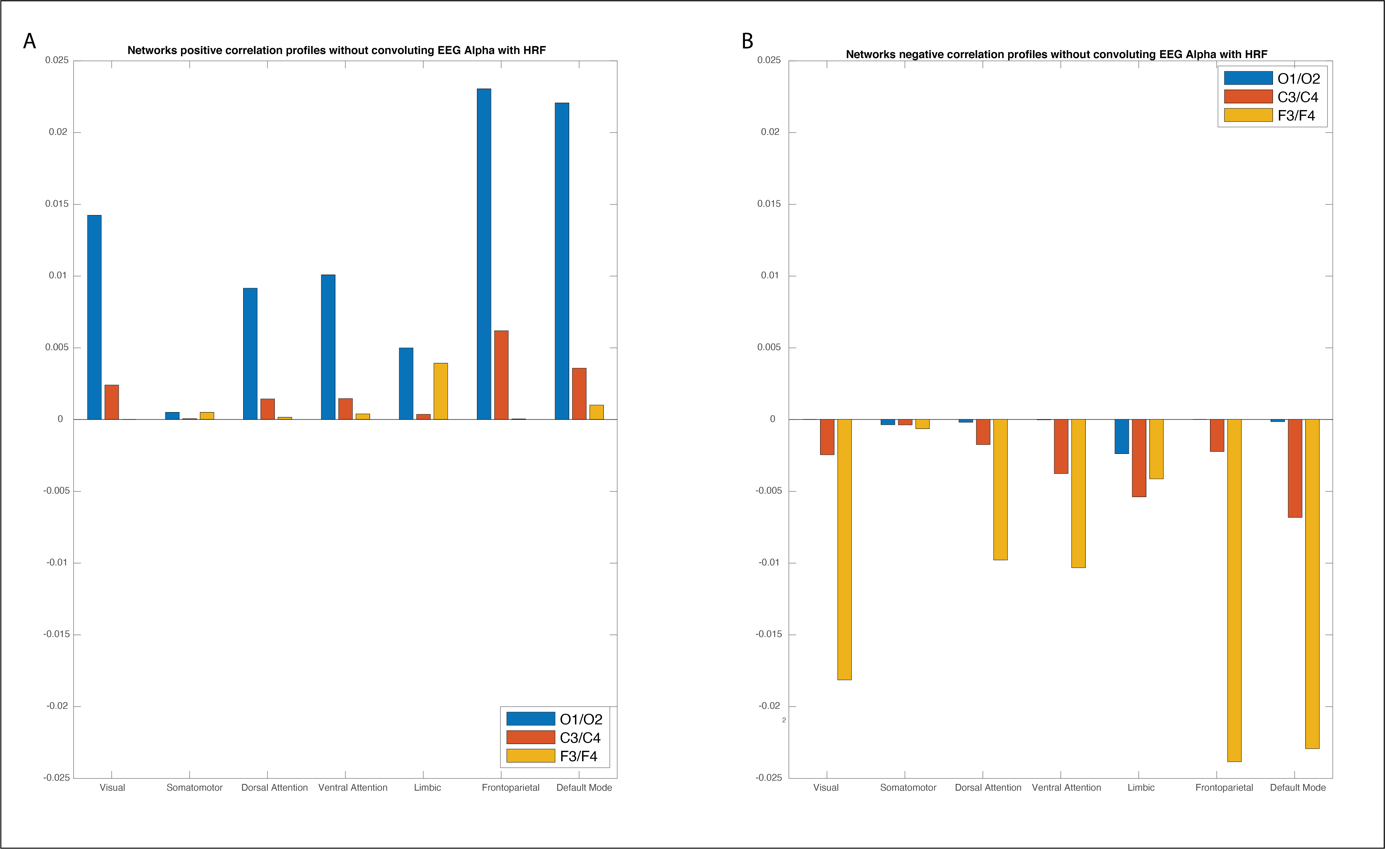

**Figure S3:** Networks correlation profiles for EEG Alpha power at different surface electrodes *without* fMRI BOLD convoluted with HRF: A) positive correlation. B) negative correlations

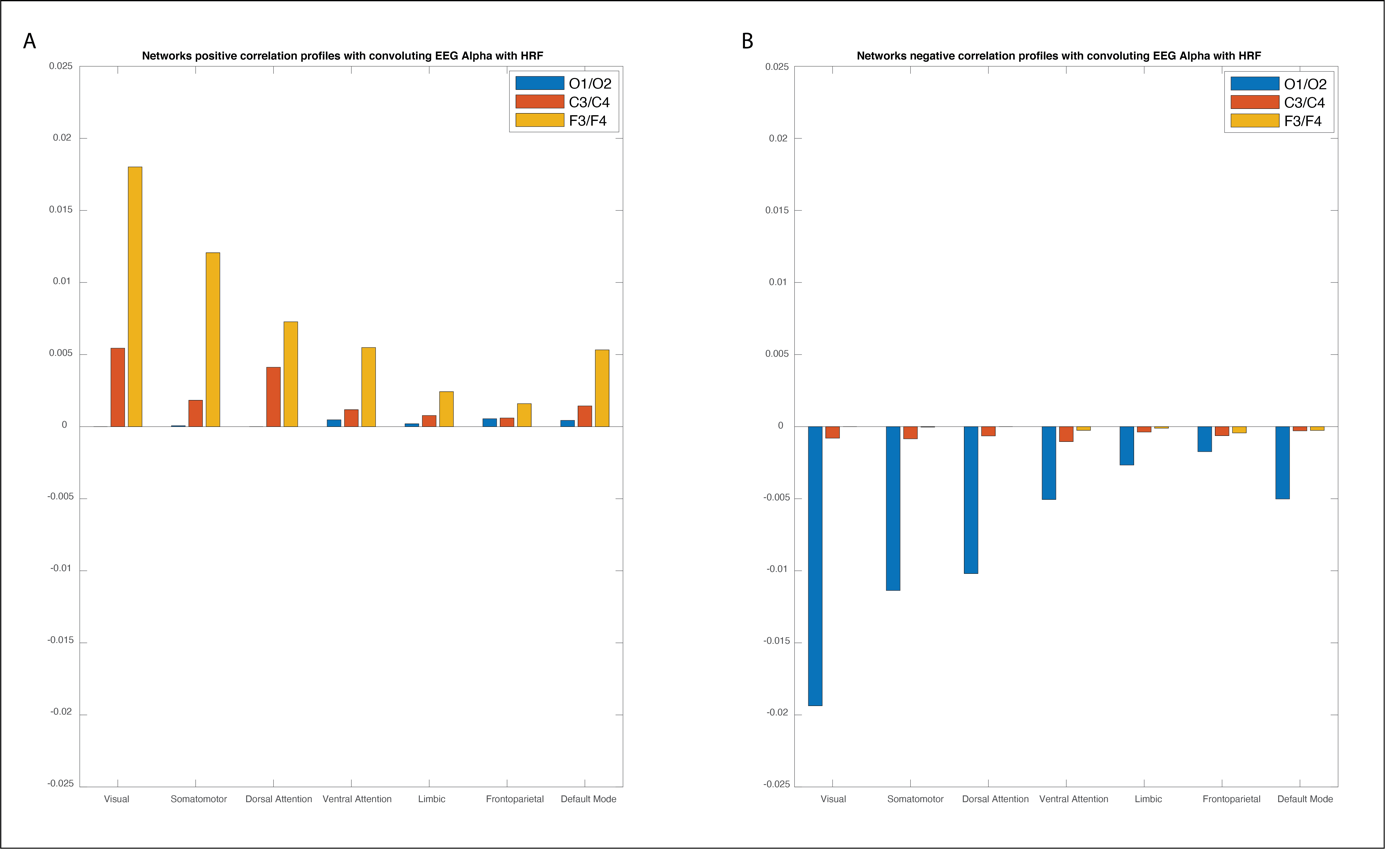

**Figure S4:** Networks correlation profiles for EEG Alpha power at different surface electrodes *with* fMRI BOLD convoluted with HRF: A) positive correlation. B) negative correlations

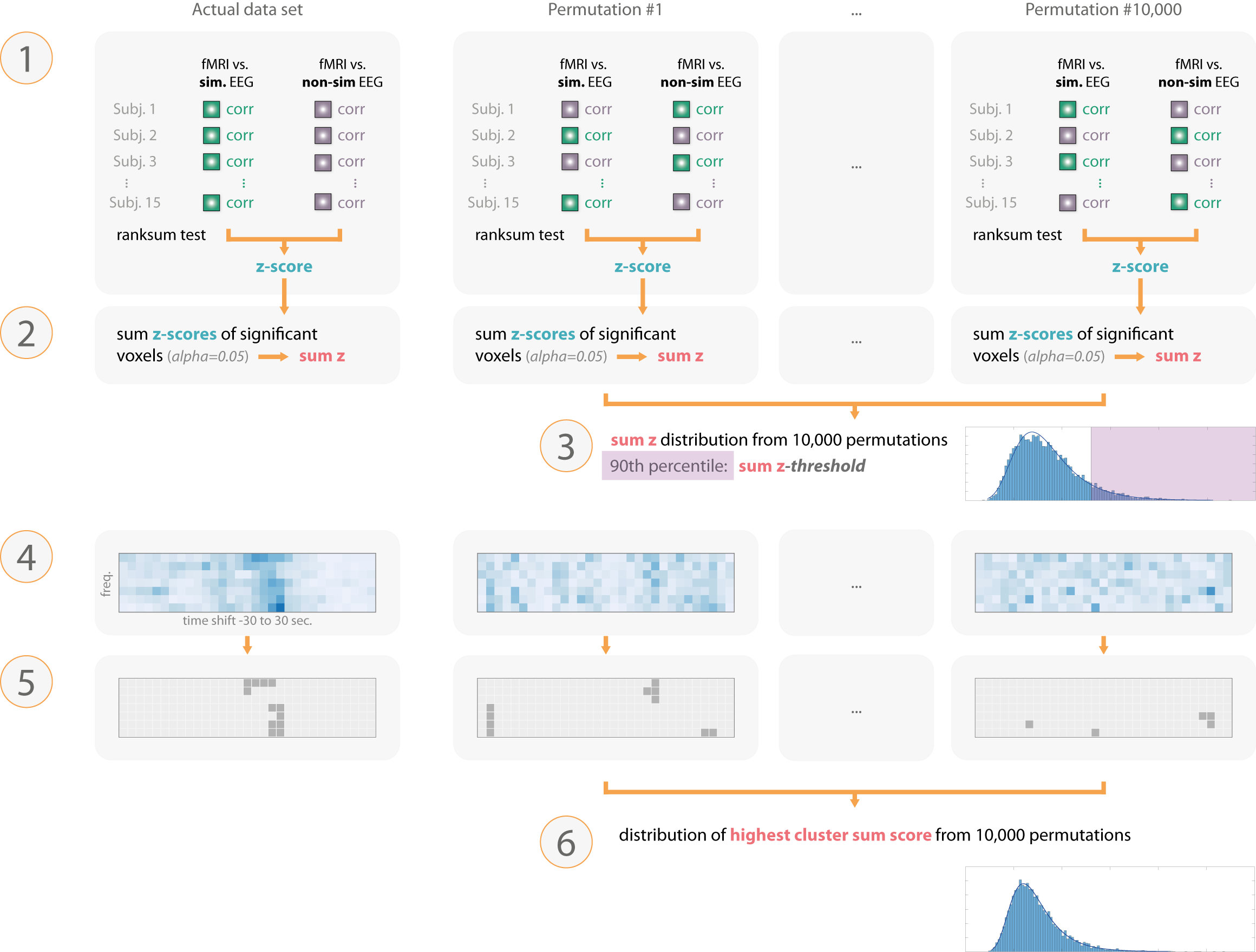

**Figure S5:** Main steps of the statistical testing.
Step 1: Time course correlations are derived from fMRI correlated with both simultaneous and non-simultaneous-EEG for each subject. The two conditions are tested at each voxel using Wilcoxon signed-rank test. The same steps were applied to each random permutation derived by randomly re-assigning the correlation values to the conditions. At each voxel a z-score was obtained.
Step 2: Significant voxels (alpha=0.05) where summed to yield “*sum z*”. This was done separately for positive and negative z-scores.

Step 3: The *sum z* value obtained from all 10,000 permutations were used to build a random distribution of the probability of obtaining a certain *sum z* by chance. A *“sum z-threshold”* was set to the 90^th^ percentile of this distribution.

Step 4: The *sum z-threshold* was used to threshold the *sum z* matrix of frequency band and time-shift combinations.

Step 5: We calculated the *cluster sum score* for each cluster found in the matrix.

Step 6: In the permuted data, the highest *cluster sum score* among the found clusters was used to derive a random distribution from all permutations. This distribution was used to calculate the probability of finding a certain *cluster sum score* by chance. Clusters derived from the observed data were tested against the distribution derived from step 6 for statistical significance. An alpha level of 0.005 (0.0025/ 20 Bonferroni corrected for multiple comparisons) was used to determine significance.

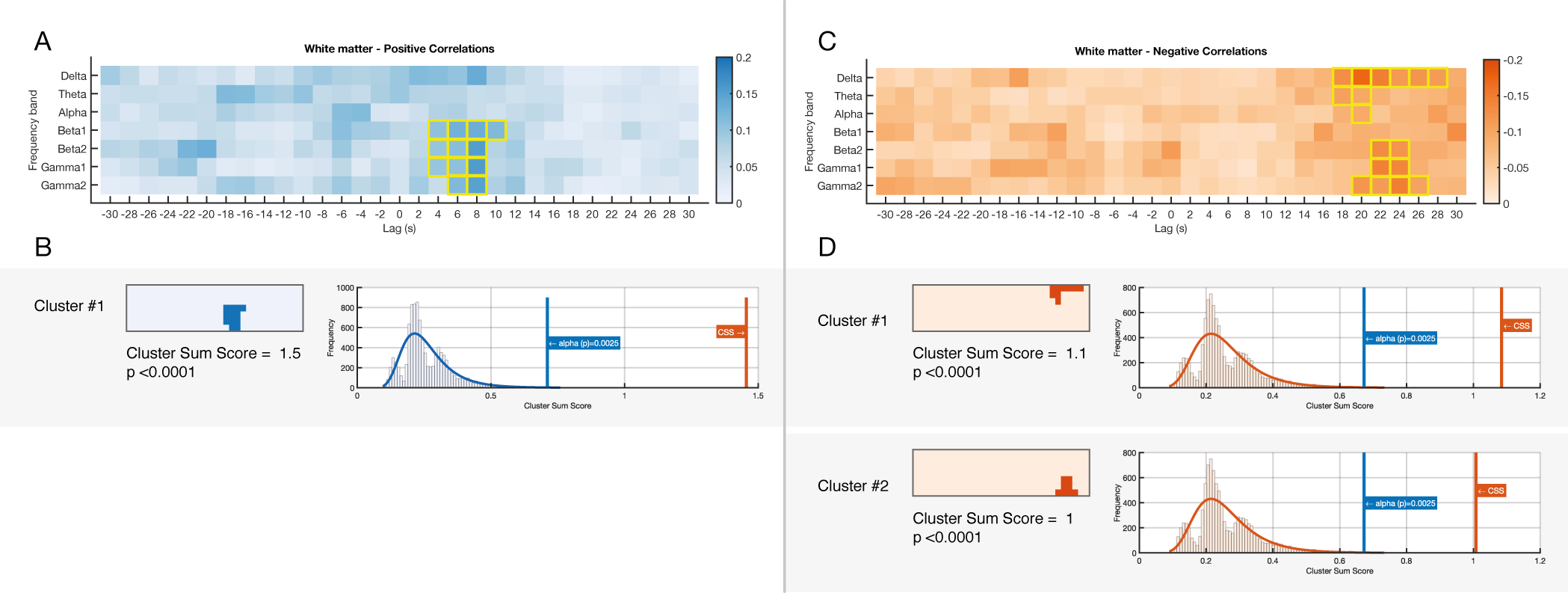

**Figure S6: EEG-fMRI significant correlations across frequencies and time-lags in white matter.**

(A) Shows a matrix of positive sum-z scores derived from significant positive correlations in the first-level statistics. Significant clusters are outlined with yellow strokes. (B) Significant clusters derived from positive correlations in A along with their *cluster sum score* compared to the random distribution. (C) Random distribution of *cluster sum scores* derived from 10,000 permutations where the two conditions. It shows the alpha level and the observed *cluster sum score* from observed data. Correlations derived from negative correlations.

D, E and F: Similar sections to A but showing results derived from significant negative correlations. For plotting purposes the color bar reflects the absolute values)

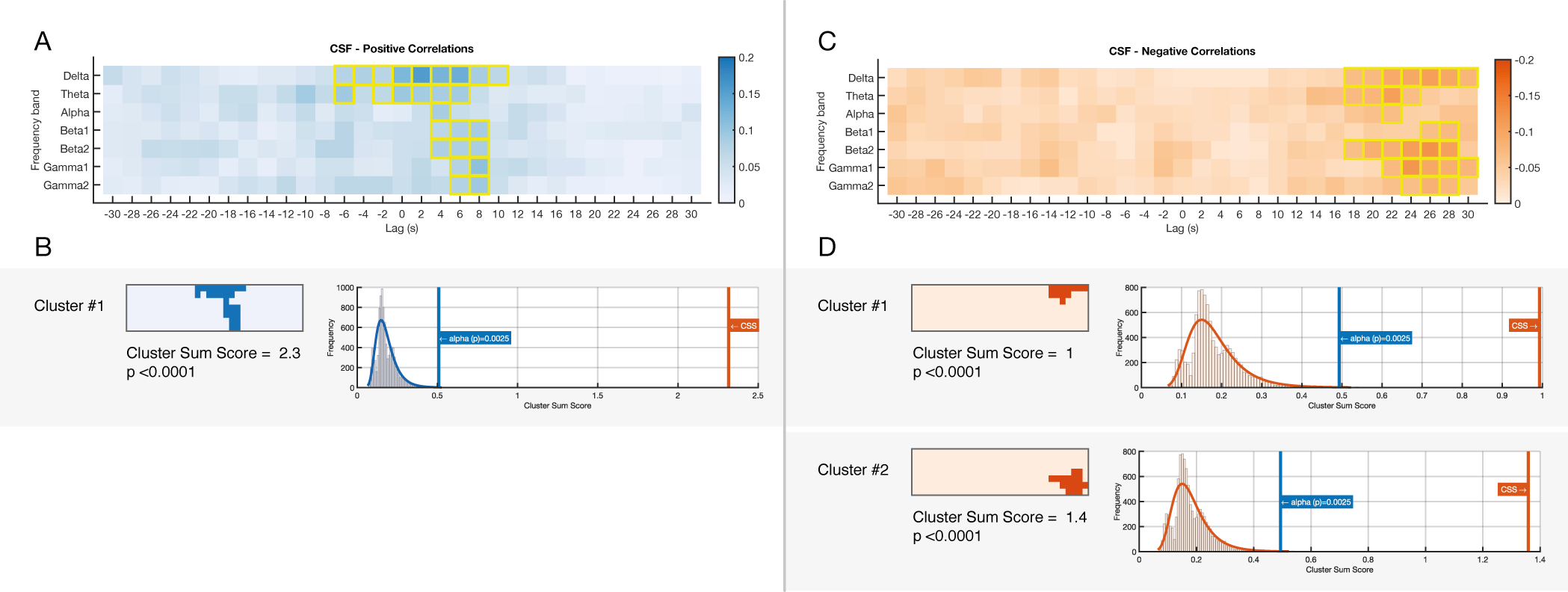

**Figure S7: EEG-fMRI significant correlations across frequencies and time-lags in CSF.**

(A) Shows a matrix of positive sum-z scores derived from significant positive correlations in the first-level statistics. Significant clusters are outlined with yellow strokes. (B) Significant clusters derived from positive correlations in A along with their *cluster sum score* compared to the random distribution. (C) Random distribution of *cluster sum scores* derived from 10,000 permutations where the two conditions. It shows the alpha level and the observed *cluster sum score* from observed data. Correlations derived from negative correlations.

D, E and F: Similar sections to A but showing results derived from significant negative correlations. For plotting purposes the color bar reflects the absolute values)

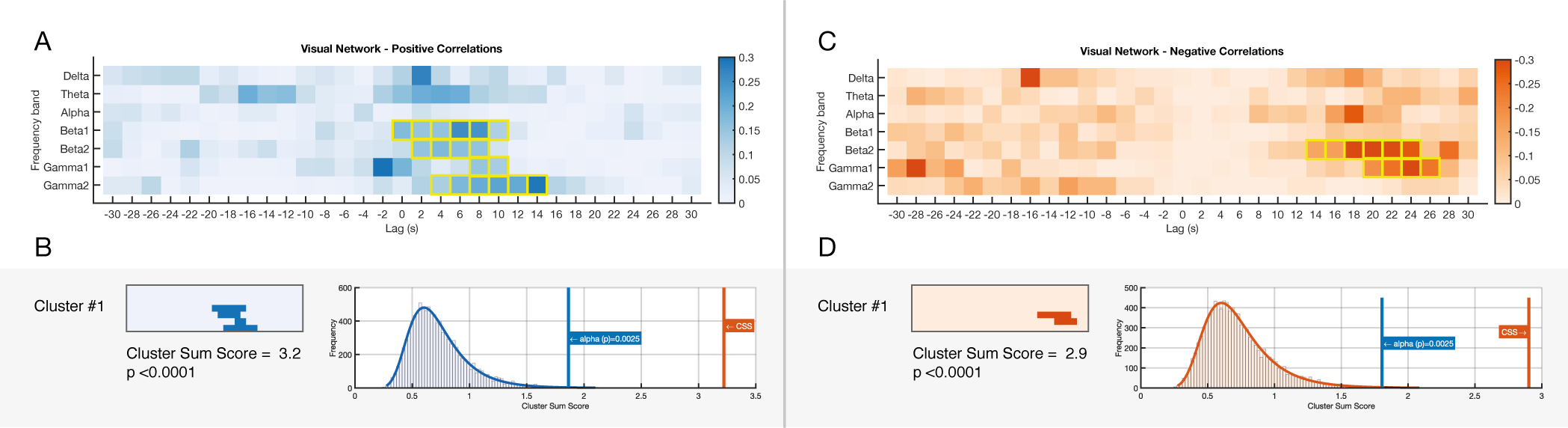

**Figure S8: EEG-fMRI significant correlations across frequencies and time-lags in the visual network.**

(A) Shows a matrix of positive sum-z scores derived from significant positive correlations in the first-level statistics. Significant clusters are outlined with yellow strokes. (B) Significant clusters derived from positive correlations in A along with their *cluster sum score* compared to the random distribution. (C) Random distribution of *cluster sum scores* derived from 10,000 permutations where the two conditions. It shows the alpha level and the observed *cluster sum score* from observed data. Correlations derived from negative correlations.

D, E and F: Similar sections to A but showing results derived from significant negative correlations. For plotting purposes the color bar reflects the absolute values)

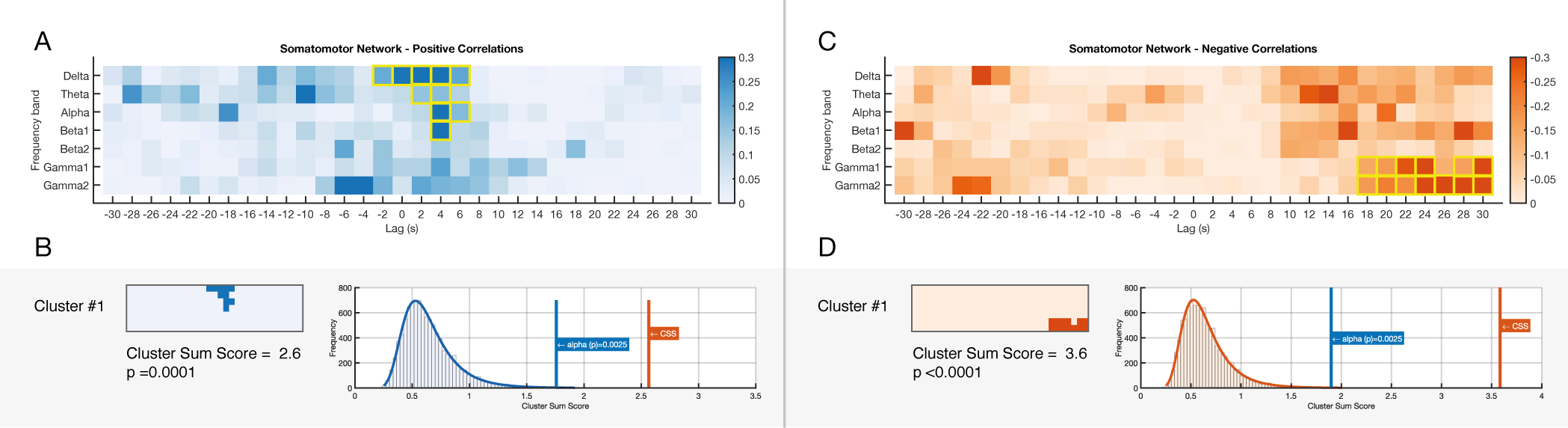

**Figure S9: EEG-fMRI significant correlations across frequencies and time-lags in the somatomotor network.**

(A) Shows a matrix of positive sum-z scores derived from significant positive correlations in the first-level statistics. Significant clusters are outlined with yellow strokes. (B) Significant clusters derived from positive correlations in A along with their *cluster sum score* compared to the random distribution. (C) Random distribution of *cluster sum scores* derived from 10,000 permutations where the two conditions. It shows the alpha level and the observed *cluster sum score* from observed data. Correlations derived from negative correlations.

D, E and F: Similar sections to A but showing results derived from significant negative correlations. For plotting purposes the color bar reflects the absolute values)

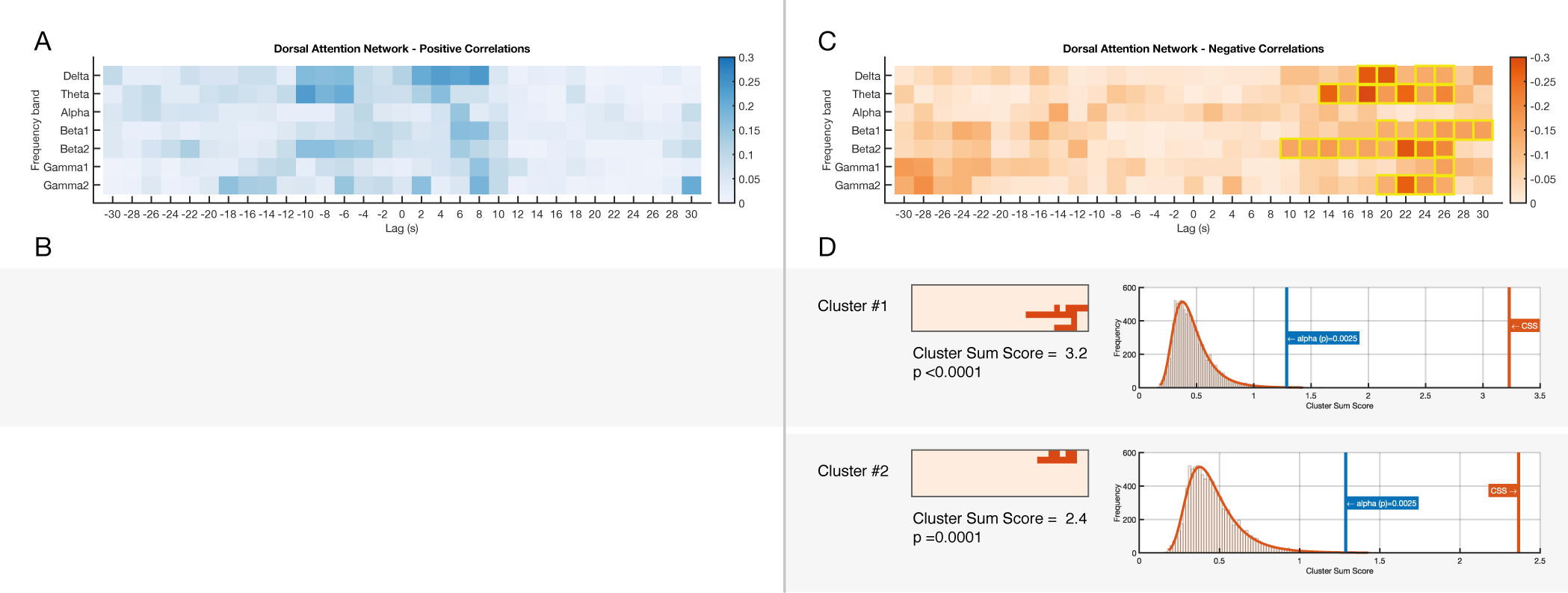

**Figure S10: EEG-fMRI significant correlations across frequencies and time-lags in the dorsal attention network.**

(A) Shows a matrix of positive sum-z scores derived from significant positive correlations in the first-level statistics. Significant clusters are outlined with yellow strokes. (B) Significant clusters derived from positive correlations in A along with their *cluster sum score* compared to the random distribution. (C) Random distribution of *cluster sum scores* derived from 10,000 permutations where the two conditions. It shows the alpha level and the observed *cluster sum score* from observed data. Correlations derived from negative correlations.

D, E and F: Similar sections to A but showing results derived from significant negative correlations. For plotting purposes the color bar reflects the absolute values)

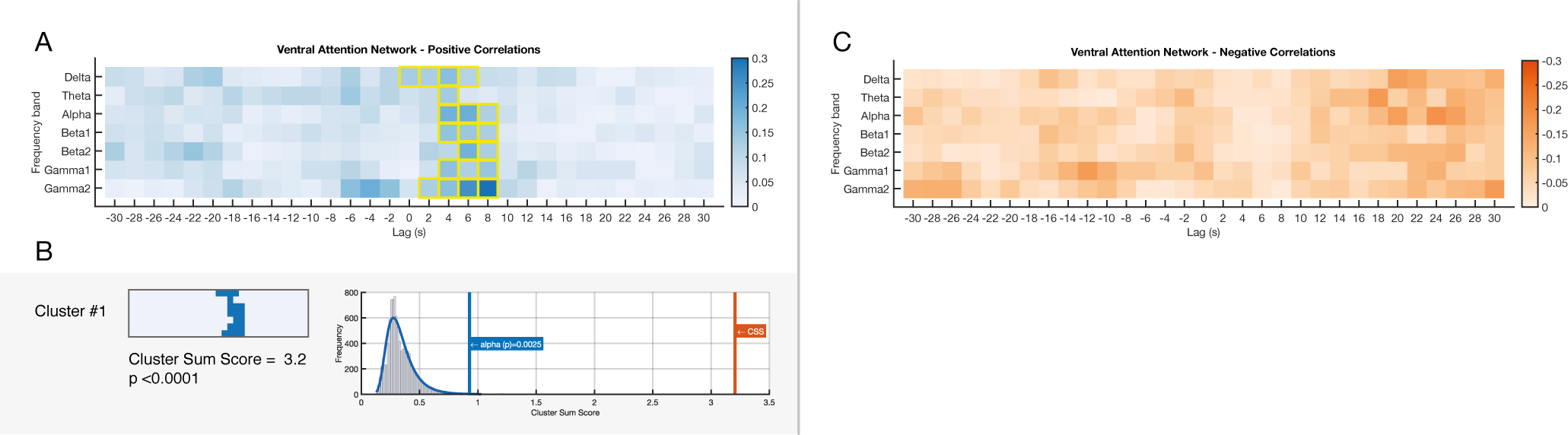

**Figure S11: EEG-fMRI significant correlations across frequencies and time-lags in the ventral attention network.**

(A) Shows a matrix of positive sum-z scores derived from significant positive correlations in the first-level statistics. Significant clusters are outlined with yellow strokes. (B) Significant clusters derived from positive correlations in A along with their *cluster sum score* compared to the random distribution. (C) Random distribution of *cluster sum scores* derived from 10,000 permutations where the two conditions. It shows the alpha level and the observed *cluster sum score* from observed data. Correlations derived from negative correlations.

D, E and F: Similar sections to A but showing results derived from significant negative correlations. For plotting purposes the color bar reflects the absolute values)

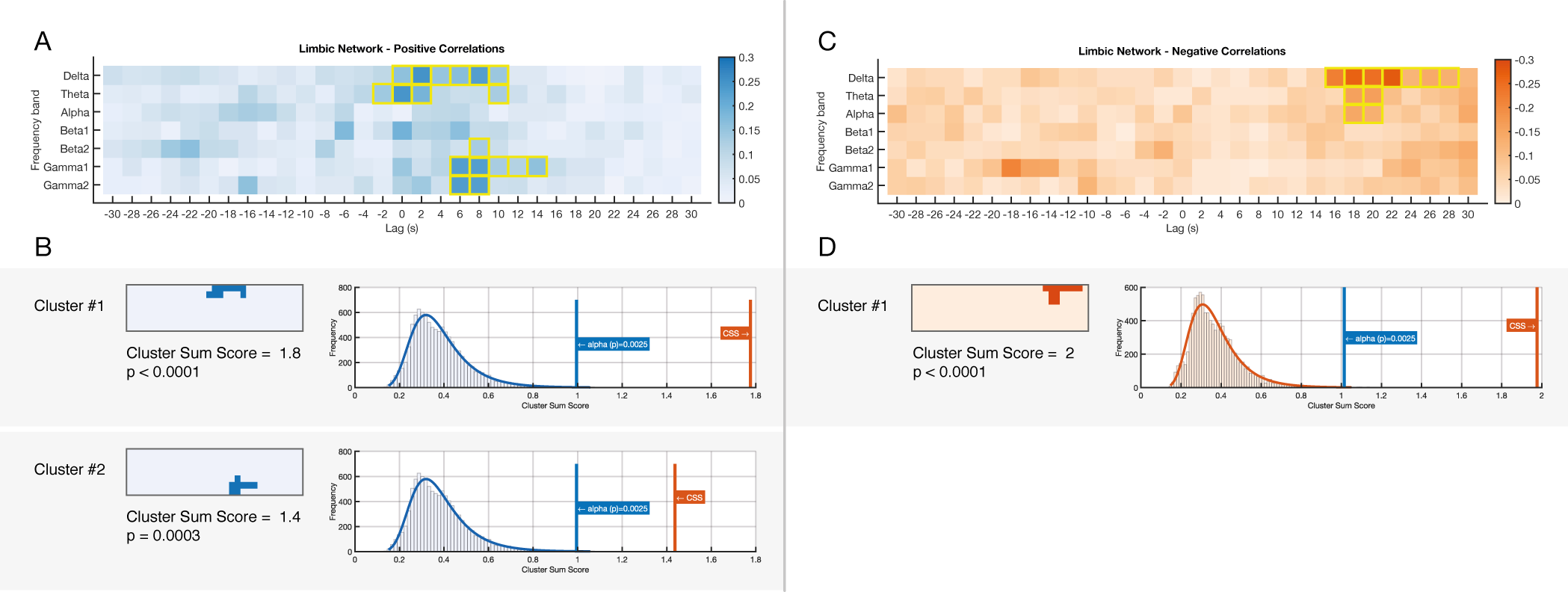

**Figure S12: EEG-fMRI significant correlations across frequencies and time-lags in the limbic network.**

(A) Shows a matrix of positive sum-z scores derived from significant positive correlations in the first-level statistics. Significant clusters are outlined with yellow strokes. (B) Significant clusters derived from positive correlations in A along with their *cluster sum score* compared to the random distribution. (C) Random distribution of *cluster sum scores* derived from 10,000 permutations where the two conditions. It shows the alpha level and the observed *cluster sum score* from observed data. Correlations derived from negative correlations.

D, E and F: Similar sections to A but showing results derived from significant negative correlations. For plotting purposes the color bar reflects the absolute values)

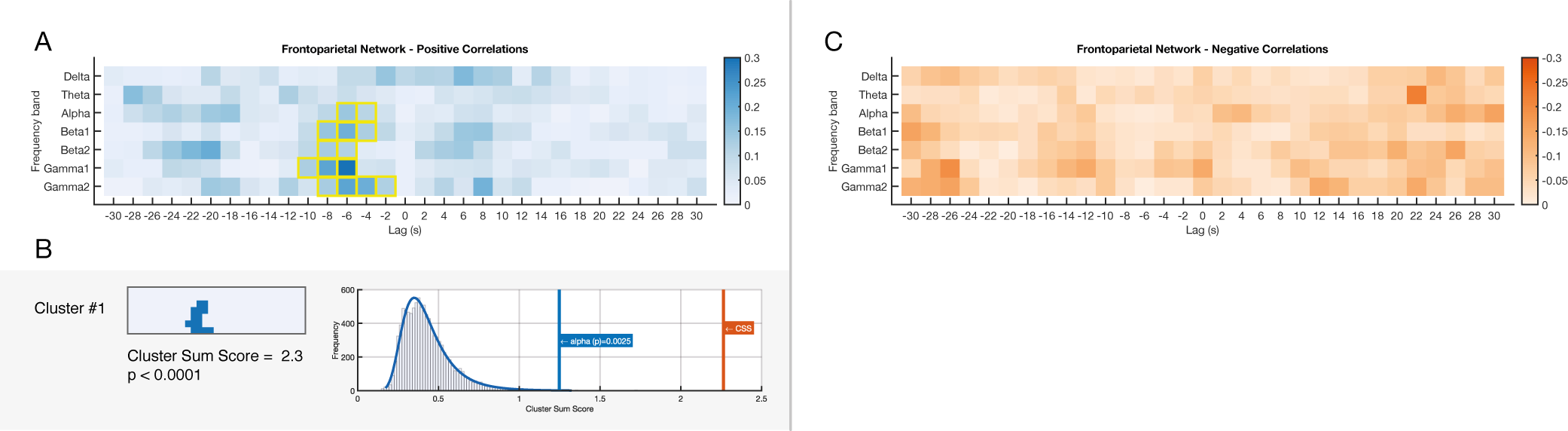

**Figure S13: EEG-fMRI significant correlations across frequencies and time-lags in the frontoparietal network.**

(A) Shows a matrix of positive sum-z scores derived from significant positive correlations in the first-level statistics. Significant clusters are outlined with yellow strokes. (B) Significant clusters derived from positive correlations in A along with their *cluster sum score* compared to the random distribution. (C) Random distribution of *cluster sum scores* derived from 10,000 permutations where the two conditions. It shows the alpha level and the observed *cluster sum score* from observed data. Correlations derived from negative correlations.

D, E and F: Similar sections to A but showing results derived from significant negative correlations. For plotting purposes the color bar reflects the absolute values)

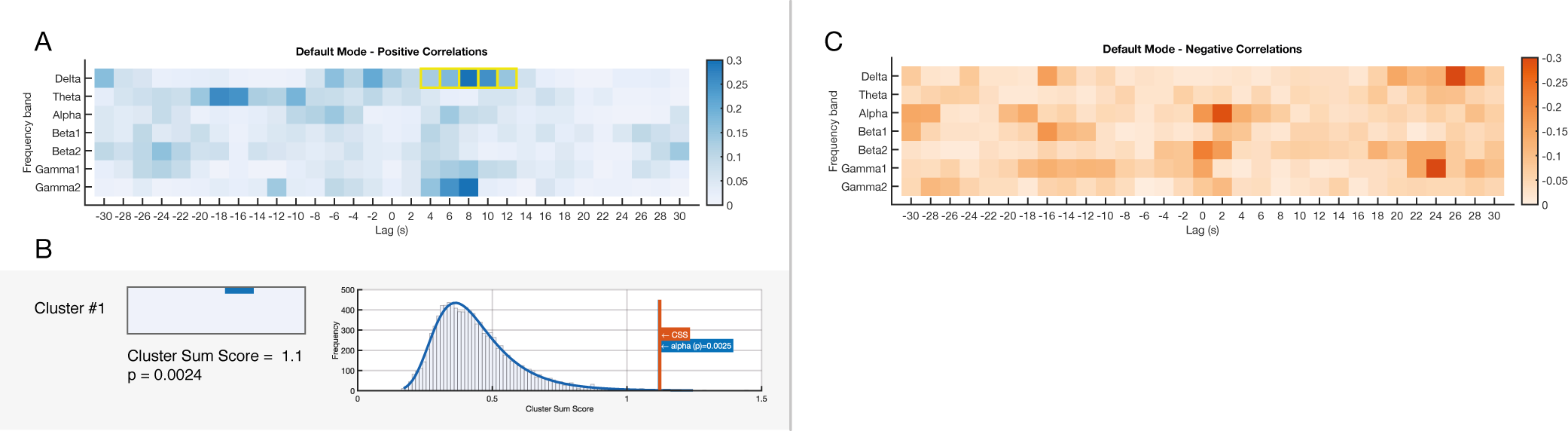

**Figure S14: EEG-fMRI significant correlations across frequencies and time-lags in the Default mode network.**

(A) Shows a matrix of positive sum-z scores derived from significant positive correlations in the first-level statistics. Significant clusters are outlined with yellow strokes. (B) Significant clusters derived from positive correlations in A along with their *cluster sum score* compared to the random distribution. (C) Random distribution of *cluster sum scores* derived from 10,000 permutations where the two conditions. It shows the alpha level and the observed *cluster sum score* from observed data. Correlations derived from negative correlations.

D, E and F: Similar sections to A but showing results derived from significant negative correlations. For plotting purposes the color bar reflects the absolute values)

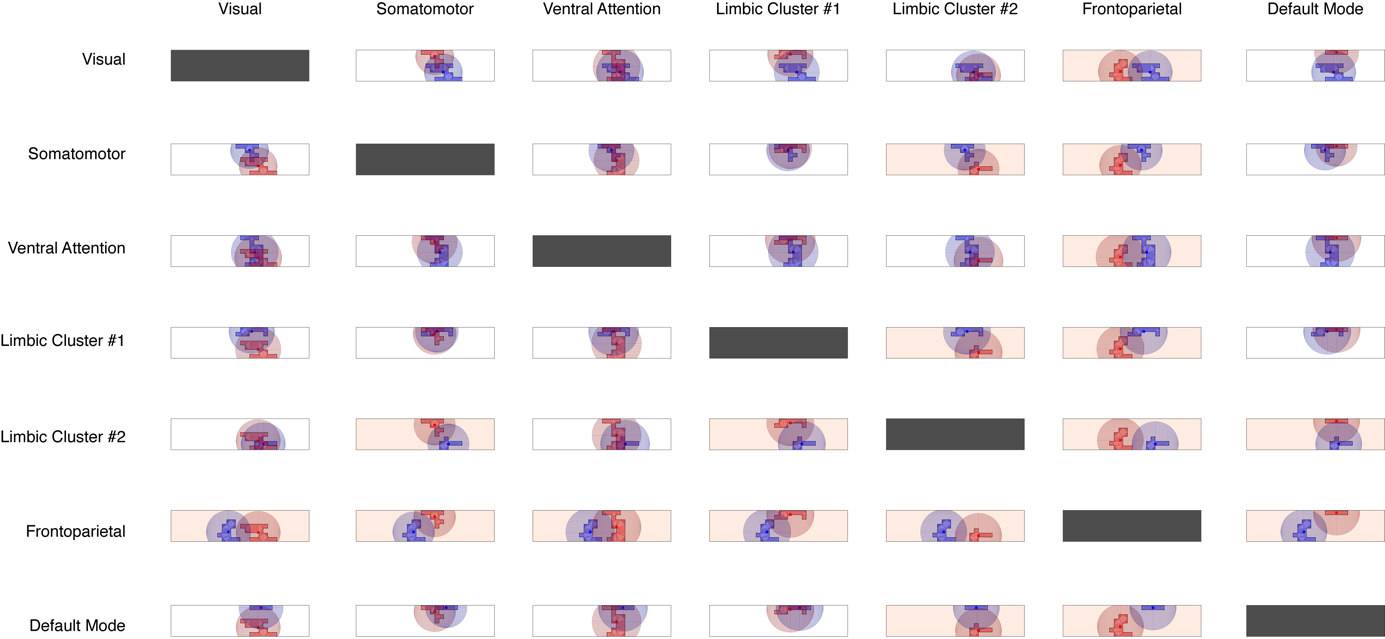

**Figure S15: Similarity tests of positive correlations clusters between networks**

For each resting state network, the cluster of significant correlations is tested against clusters derived from other networks. The distance between the centroids of the two clusters is tested against the distribution formed by the distance of all the points belonging to each cluster and its centroid. Red background denotes that the patterns in the two networks are significantly different (alpha = 0.05).

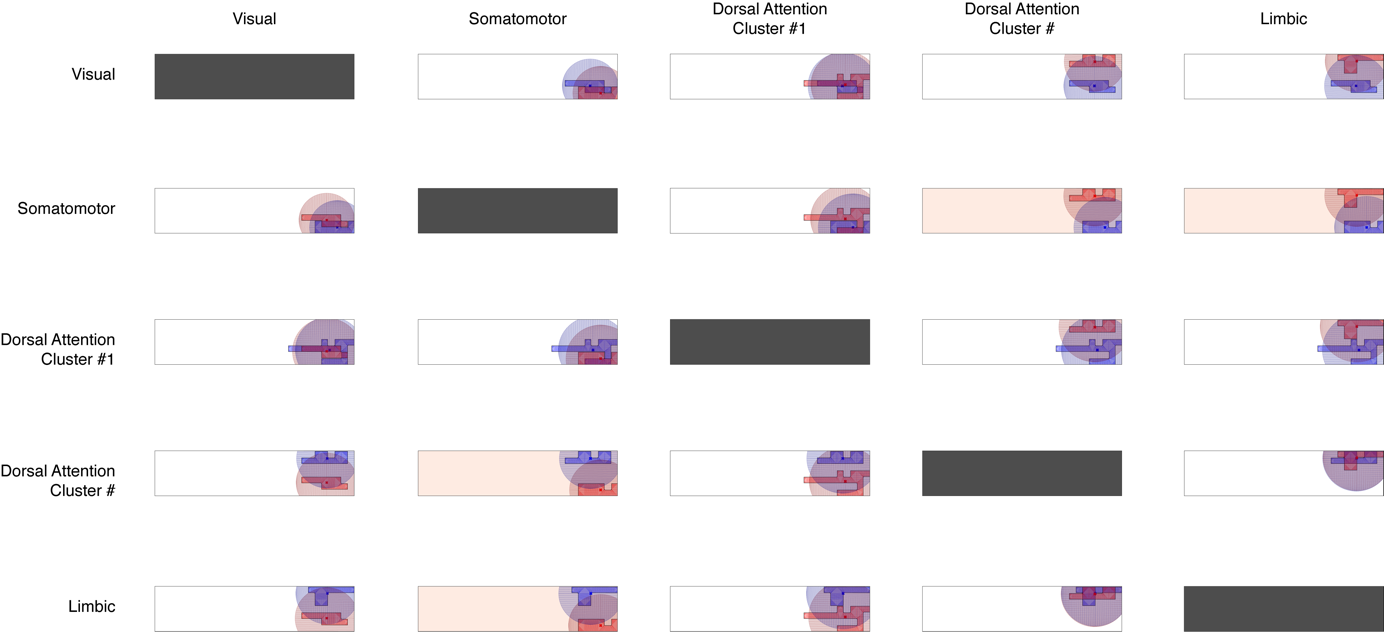

**Figure S16: Similarity tests of negative correlations clusters between networks**

For each resting state network, the cluster of significant correlations is tested against clusters derived from other networks. The distance between the centroids of the two clusters is tested against the distribution formed by the distance of all the points belonging to each cluster and its centroid. Red background denotes that the patterns in the two networks are significantly different (alpha = 0.05).
